# Cilevirus and dichorhavirus glycoproteins target overlapping host proteins in the *Brevipalpus yothersi* vector

**DOI:** 10.64898/2026.07.17.739234

**Authors:** C. Chabi-Jesus, P.L. Ramos-González, T.C. Bernardino, L.G.O. Guardalini, S.A.C. Jorge, A.D Tassi, R. Harakava, E.W. Kitajima, A. Whitfield, J. Freitas-Astúa

## Abstract

Enveloped plant viruses are rare, and accumulating evidence suggests deep evolutionary ties to arthropod hosts. *Brevipalpus-*transmitted viruses (BTVs), which include cileviruses, higreviruses, and dichorhaviruses, are transmitted exclusively by *Brevipalpus* mites, causing localized infections in plants. Some cileviruses and dichorhaviruses can also replicate within vector cells. Despite their enveloped virions, the involvement of virion structural proteins mediating the interaction with the vector remains unresolved. Here, we investigated the putative glycoproteins P61 of citrus leprosis virus C (CiLV-C; *Cilevirus, Kitaviridae*) and G of clerodendrum chlorotic spot virus (ClCSV; *Dichorhavirus, Rhabdoviridae*) to define their interaction networks in the *Brevipalpus yothersi* vector. Using membrane-based yeast two-hybrid screening with a *B. yothersi* cDNA library, we identified 73 interactors for P61 and 162 for G, including a shared core set enriched for membrane-associated proteins involved in intracellular trafficking, ER–Golgi dynamics, protein synthesis and folding, and signal transduction. Numerous hypothetical membrane proteins emerged as strong candidates for viral receptors or co-receptors. Three proteins (ARF1, SERP2, and a hypothetical transmembrane protein) were validated by BiFC and Co-IP in *Spodoptera frugiperda* (Sf9) cells, confirming interactions with both viral proteins in an arthropod-like environment. Together, this work provides the first comprehensive interactome of BTV glyco-like proteins with those of its natural vector and reveals partially convergent host-interaction strategies between phylogenetically distinct viruses, establishing a mechanistic basis for dissecting BTV transmission.

**IMPORTANCE:** BTV’s damage economically important crops worldwide, and citrus leprosis (CL) is a significant disease in the Brazilian citrus belt. Brazil is responsible for over 75% of global sweet orange juice production and widely cited estimates indicate that CL causes >US$69 million in annual yield losses and acaricide-based control of *Brevipalpus yothersi*, the vector of CiLV-C and several other BTVs. Despite this impact, the molecular basis of BTV acquisition, persistence, and transmission by *Brevipalpus* spp. remains largely unexplored. Here, we provide the first systematic map of mite proteins interacting with glycoproteins from two BTVs belonging to distinct viral families. These interactomes reveal key components of the cellular machinery targeted by cileviruses and dichorhaviruses in their arthropod vector, offering mechanistic insight into how these viruses establish and maintain infections in mites. This work establishes a foundational framework for future studies aimed at disrupting BTV transmission and developing sustainable control strategies.

## INTRODUCTION

*Brevipalpus* mites (Acari: Tenuipalpidae) are cosmopolitan, highly polyphagous arthropods, commonly known as false spider mites (1,2). Although the genus comprises more than 300 described species, fewer than ten are recognized as economically important pests and confirmed vectors of plant-infecting viruses (1,3). Known vector mites can colonize more than 900 plant species, including citrus, coffee, passionfruit, papaya, tea, grapes, and other major crops cultivated in tropical and subtropical regions (4,5).

As virus vectors, *Brevipalpus* mites transmit a unique assemblage of phylogenetically unrelated RNA viruses collectively known as *Brevipalpus*-transmitted viruses (BTVs) (6–8). BTVs span three viral genera: *Cilevirus* and *Higrevirus* (family *Kitaviridae*; positive-sense RNA) and *Dichorhavirus* (family *Rhabdoviridae*; negative-sense RNA) (7–9). Among BTV-associated diseases, citrus leprosis (CL) is the most devastating. In Brazil’s main citrus belt, located in the States of São Paulo and Minas Gerais, CL requires more than US$54 million annually for acaricide-based vector control and accounts for over 19% of disease-associated premature fruit drop, resulting in losses exceeding US$15.6 million per year (10,11). Although multiple BTV have been detected in CL-affected trees, citrus leprosis virus C (CiLV-C; *Cilevirus leprosis*) is the predominant causal agent across the Americas (12,13) and is transmitted mainly by *Brevipalpus yothersi*, with experimental reports of transmission by *B. papayensis* (14,15).

Despite their phylogenetic divergence, many BTVs share several hallmark traits: production of bacilliform-like virions, localized non-systemic lesions in plants, putative glycoproteins, and phylogenetic affinities with arthropod-infecting viruses (6–8). Together, these features support the hypothesis that BTVs originated from arthropod-associated ancestors and subsequently adapted to infect plant tissues (6,8). Nonetheless, the molecular basis underlying their specificity for *Brevipalpus* vectors remains poorly understood.

For CiLV-C, virions and antigenomic RNA have been detected in viruliferous *B. yothersi*, and antigenomic RNA may persist at almost the same level for at least 15 days in mites removed from infected plants (8). These observations are consistent with a propagative mode of transmission (8,16,17). Strong evidence supports dichorhavirus replication within mite tissues; their viral titers increase after acquisition, viroplasms are present in the anterior and midgut and in salivary glands, and long latent periods are required for transmission (18–21). Furthermore, dichorhaviruses belong to the family *Rhabdoviridae* and are closely related phylogenetically to viruses that replicate in both arthropod and plant hosts (19,21). Clerodendrum chlorotic spot virus (ClCSV; *Dichorhavirus clerodendri*), for example, is detectable in *B. yothersi* (22) and becomes transmissible ≥20 days after acquisition (A.D.Tassi, unpublished data).

Given that many BTVs produce enveloped virions, their glycoproteins are presumed to mediate vector interaction and cellular entry (6,23). In CiLV-C, P61 is a strong glycoprotein candidate based on predicted structural features and the ability to induce ER stress, unfolded protein response activation, and hypersensitive-like symptoms in *Nicotiana benthamiana* (8,24,25). Conversely, dichorhaviruses are traditionally considered non-enveloped viruses (7,26). However, they encode a G protein with sequence and functional parallels to rhabdoviridal glycoproteins (27,28), and natural variation in G correlates with Brevipalpus species groups (29), suggesting a role in vector specificity. Moreover, studies of enveloped plant RNA viruses have shown that viral glycoproteins can be dispensable for infection in plants while remaining essential for infection and transmission by their arthropod vectors (30).

In this study, we investigate the molecular interface between BTV glycoproteins and their mite vector using P61 (CiLV-C) and G (ClCSV) as model systems. We performed membrane-based yeast two-hybrid (MbY2H) screenings with a B. yothersi cDNA library to map protein–protein interactions and validated representative candidates in Spodoptera frugiperda (Sf9) cells using bimolecular fluorescence complementation (BiFC) and co-immunoprecipitation (Co-IP). Our findings reveal conserved and virus-specific vector components targeted by BTV glycoproteins and identify key mite proteins that may play roles in virus acquisition, persistence, and transmission.

## MATERIALS AND METHODS

### Virus isolates and *Brevipalpus* vector colonies

The putative glycoprotein P61 of CiLV-C strain SJP and the glycoprotein G of ClCSV were selected as representatives of cileviruses and dichorhaviruses, respectively. CiLV-C SJP was obtained from naturally infected *Citrus sinensis* plants collected in Jaboticabal, São Paulo, Brazil (2020), whereas ClCSV was sourced from symptomatic *Clerodendrum* sp. collected in Piracicaba, São Paulo (2021).

Non-viruliferous *Brevipalpus yothersi* colonies were maintained on *Citrus latifolia* fruits at the Instituto Biológico, São Paulo, Brazil. A second colony reared on common beans (*Phaseolus vulgaris*) leaves was provided by Dr. Aline Tassi (University of Florida, Homestead, USA). All colonies were maintained under controlled environmental conditions (25°C, 16 h light/8 h dark, 50–60% relative humidity). The virus-free status of the mite colony was routinely verified by RT-PCR.

### Construction, expression, and validation of MbY2H bait plasmids

Signal peptide (SP) and transmembrane (TM) domain predictions (SignalP v6.0, DeepTMHMM) (31,32) indicated that both viral proteins are type I membrane proteins with N-terminal SPs and C-terminal TM regions. To promote correct membrane insertion in yeast, coding sequences lacking the predicted SPs were amplified using PrimeSTAR® GXL Premix (Takara Bio) with gene-specific primers (Supplementary Table S1).

PCR amplicons were gel-purified and assembled into the linearized pBTR3-SUC vector (Dualsystems Biotech®) via HiFi DNA Assembly (NEB), yielding pBTR3-SUC:P61-CiLV-C and pBTR3-SUC:G-ClCSV. Constructs were transformed into *Escherichia coli* DH10β, selected on LB-kanamycin, screened by colony PCR, and fully sequence-verified by Oxford Nanopore long-read sequencing (Plasmidsaurus).

Functional tests followed the manufacturer’s protocol (Takara Bio). Bait constructs were transformed into *Saccharomyces cerevisiae* NMY51 together with pOst1-NubI (positive control) to assess correct membrane localization and reporter activation. Autoactivation was evaluated by co-transforming bait plasmids with an empty prey vector (pPR3-N) and plating on QDO medium supplemented with increasing concentrations of 3-AT (0, 1, 5, 10 mM). Growth was recorded after 72 h.

### RNA extraction and ds-cDNA synthesis from *Brevipalpus yothersi*

RNA extraction from *B. yothersi* is challenging due to the small body size (adults measure 0.25–0.30 mm in length), low RNA content, and chitin-rich cuticle. Because high-quality, high-yield RNA was essential for cDNA library construction, two extraction kits and two RNA-concentration strategies were tested in parallel to maximize the likelihood of obtaining suitable material: RNAqueous™ (Thermo Fisher Scientific) and Quick-RNA Tissue/Insect Microprep (Zymo Research). Approximately 1,000 mites (lime-fed colony) and 2,000 mites (bean-fed colony) were processed. When required, RNA was concentrated either by ethanol precipitation (0.1 vol of 3 M sodium acetate, pH 5.5) or using the Monarch® RNA Cleanup Kit (NEB). RNA quality and integrity were assessed with the Agilent TapeStation 4200. Full-length cDNA was synthesized using the SMART® cDNA Library Construction Kit (Takara Bio) with 2 µg of total RNA, followed by LD-PCR amplification with the Advantage® 2 kit. LD-PCR products were checked on 1.2% agarose gels to assess size distribution.

### Construction of the pPR3-N prey cDNA library

To enrich ORFs compatible with MbY2H screening, LD-PCR products were digested with SfiI and separated on 1.2% agarose gels. DNA fragments between 1–2 kb were excised and purified with the Monarch® DNA Gel Extraction Kit (NEB). Plasmid vector preparation consisted of SfiI digestion (16 h at 16 °C) of pPR3-N, followed by gel purification to remove undigested plasmid. Ligations were performed using 500 ng of linearized pPR3-N and graded insert volumes (0.5–1.5 µL). Reaction mixtures were electroporated into *E. coli* DH10β, recovered for 30 min in SOC medium, and plated onto 150-mm LB–ampicillin plates to maximize coverage.

After overnight growth at 37 °C, colonies from all plates were scraped using sterile microscope slides, pooled into 3 L of LB medium, incubated for 3 h at 30 °C (250 rpm), and harvested by centrifugation (4,000×g, 10 min). Plasmid DNA was extracted using the QIAGEN Plasmid Maxi Kit (Qiagen). Insert size distribution and library complexity were estimated by colony PCR of randomly selected clones using universal pPR3-N primers.

### MbY2H screening for interactors of P61-CiLV-C and G-ClCSV

MbY2H screening was conducted in *S. cerevisiae* NMY51 using the LiAc/SS-DNA/PEG transformation protocol. Yeasts were co-transformed with the prey library and either pBTR3-SUC:P61 (cilevirus protein) or pBTR3-SUC:G (dichorhavirus protein). After recovery, transformation mixtures were plated onto 150-mm QDO plates containing 1 mM 3-AT (P61 screen) or 5 mM 3-AT (G screen). Plates were incubated for four days at 30 °C.

Colonies growing under selection were picked, and prey plasmids were recovered using the Zymoprep Yeast Miniprep Kit (Zymo Research). Prey plasmids were electroporated into *E. coli* DH5α, and two colonies per yeast isolate were screened by PCR. Inserts ≥0.9 kb were purified and submitted for Sanger sequencing.

### Bioinformatic analysis of prey interactors

Redundant sequences were removed, and unique ORFs were annotated by BLASTn and BLASTx searches against the NCBI nr/nt database (33) and the *B. yothersi* genome (34) and transcriptome (35). Additional comparisons were performed with *B. papayensis*, *B. californicus* (33,34), and *B. obovatus* (36). For poorly annotated proteins, homologs from *Tetranychus urticae* (37)were used to assist functional inference.

Functional categorization was preformed using UniProt (38) and eggNOG-mapper (39). Subcellular localization predictions were obtained using BUSCA (40), integrating SignalP v6.0 (31), DeepTMHMM (32), GPI-anchor prediction, and other modules. DeepGOPlus (41), HHblits, HHpred, and HMMER (42) were used for remote homology detection of hypothetical proteins.

### Individual MbY2H assays and interaction stringency tests

Candidate proteins, including predicted transmembrane and membrane-associated proteins, were retested with P61 and G in individual MbY2H assays (Supplementary Table S2). Interactions were scored based on HIS⁺/ADE⁺/lacZ⁺ phenotypes on QDO medium containing 40 µg/mL X-Gal.

To assess stringency, interactions were evaluated under increasing 3-AT concentrations (Supplementary Table S2). For P61, plates contained 0, 2.5, or 5 mM 3-AT; for G, 0, 5, or 10 mM 3-AT. Overnight cultures grown in QDO medium were normalized to OD₅₆₀ = 0.8, and 16 µL drops were spotted in triplicate. Growth was scored visually from 0 (no growth) to 3 (strong growth) after 72 h. Negative controls included bait + empty prey vector; positive controls included pOst1-NubI and the pTSU2-APP × pNubG-Fe65 interaction pair.

### Prediction of 3D structures and interaction modeling

Three *B. yothersi* proteins (ARF1, SERP2, and bryot216g00140) were selected for *in silico* structural analysis due to consistent MbY2H interactions with both viral glycoproteins. Signal peptides and TM helices were predicted using SignalP 6.0(31) and DeepTMHMM(43); PTMs were assessed with MusiteDeep(44) and NetNGlyc (45). AlphaFold 3.0 was used to generate structural models and interaction predictions. Signal peptide and transmembrane domain regions were removed before modeling to avoid artifacts in docking predictions (Supplementary Table S3). Model reliability was assessed using pLDDT, PAE, and interface scoring.

### Plasmid construction for heterologous expression, BiFC, and Co-IP

Coding sequences for ARF1, SERP2, and bryot216g00140 were amplified from *B. yothersi* cDNA (Supplementary Table S1) and cloned into modified pI-RFP (46) vectors for subcellular localization in Sf9 cells. Viral P61 and G coding sequences were similarly cloned into pI-GFP (46).

For BiFC assays, the viral glycoproteins were fused at their C-terminus to the C-terminal half of YFP (cYFP), followed by a 6×FLAG tag (Supplementary Figure S5). Mite proteins were fused to the N-terminus or C-terminus of the N-terminal half of YFP (nYFP) and a 6×Myc tag. Both orientations were constructed and empirically evaluated, and the configuration yielding optimal expression and BiFC signal was used in subsequent assays.

To allow co-expression of both partners from a single vector, dual-promoter plasmids were engineered by introducing a second expression cassette downstream of the existing transcription unit within the pIcYFP backbone (Supplementary Figure S5). This strategy generated dual-promoter plasmids capable of simultaneously expressing glycoprotein– cYFP–FLAG and mite–nYFP–Myc fusions in Sf9 cells. An MBP-based BiFC positive control was also constructed. All plasmids were validated by Nanopore sequencing.

### Heterologous expression, localization, and BiFC in Sf9 cells

Sf9 cells (0.5 × 10⁶ per well) were transfected with 2 µg plasmid DNA using Cellfectin® or Lipofectamine™ Stem (Thermo Fisher Scientific). After 5 h at 22 °C, the medium was replaced, and cultures were incubated at 27 °C. Fluorescence was monitored at 6, 12, 24, 48, 72, 96 and 120 hours post-transfection (hpt) using an Agilent BioTek microscope.

Cells were transferred to glass-bottom dishes for confocal imaging (Zeiss LSM 980). DAPI, ER-Tracker Blue-White DPX (Thermo Fisher Scientific), and BODIPY TR-Ceramide (Thermo Fisher Scientific) were used to visualize nuclei, ER, and Golgi compartments, respectively. BiFC complexes were evaluated for intensity and spatial patterns compatible with the localization of each partner protein.

### Co-immunoprecipitation and Western blotting

Co-IP assays were performed at 24 hpt for P61 and 72 hpt for G, using 2 × 10⁶ and 1 × 10⁶ transfected Sf9 cells, respectively, corresponding to their peak expression times. Cells were lysed in Pierce™ IP Lysis Buffer, clarified, and incubated with anti-FLAG affinity gel (Selleckchem). The resin was 3x washed with PBS + 0.1% Tween-20, and bound complexes were eluted in Laemmli buffer and boiled for 10 min.

Proteins were resolved by SDS-PAGE, transferred to PVDF membranes, and blocked in 5% milk/TBST. Prey proteins were detected with anti-Myc (9E10; Thermo Fisher Scientific) followed by TrueBlot® ULTRA HRP secondary antibody (Rockland) to avoid IgG interference. Viral proteins were detected using anti-FLAG–HRP. Signals were visualized with an iBright™ FL1500 system.

## RESULTS

### P61 of CiLV-C and G of ClCSV are representative candidate glycoproteins of *Brevipalpus*-transmitted viruses

To identify representative candidate glycoproteins for investigating virus-vector interactions in cileviruses and dichorhaviruses, we first analyzed the predicted structural features of P61-CiLV-C and G-ClCSV, respectively. Both proteins displayed the characteristic architecture of membrane-associated viral glycoproteins, including an N-terminal signal peptide, C-terminal transmembrane domain(s), and predicted glycosylation sites (Figure 1). P61 was predicted to contain two transmembrane domains (residues 462–484 and 489–511) and two high-confidence N-glycosylation sites (N22 and N323), whereas G contained one transmembrane domain (residues 482–504), three high-confidence N-glycosylation sites (N231, N351, and N387), and two predicted O-glycosylation sites (O101 and O368). Although the glycoprotein nature of these proteins remains to be experimentally validated, their predicted structural organization is consistent with that of candidate viral glycoproteins.

**Figure 1.**
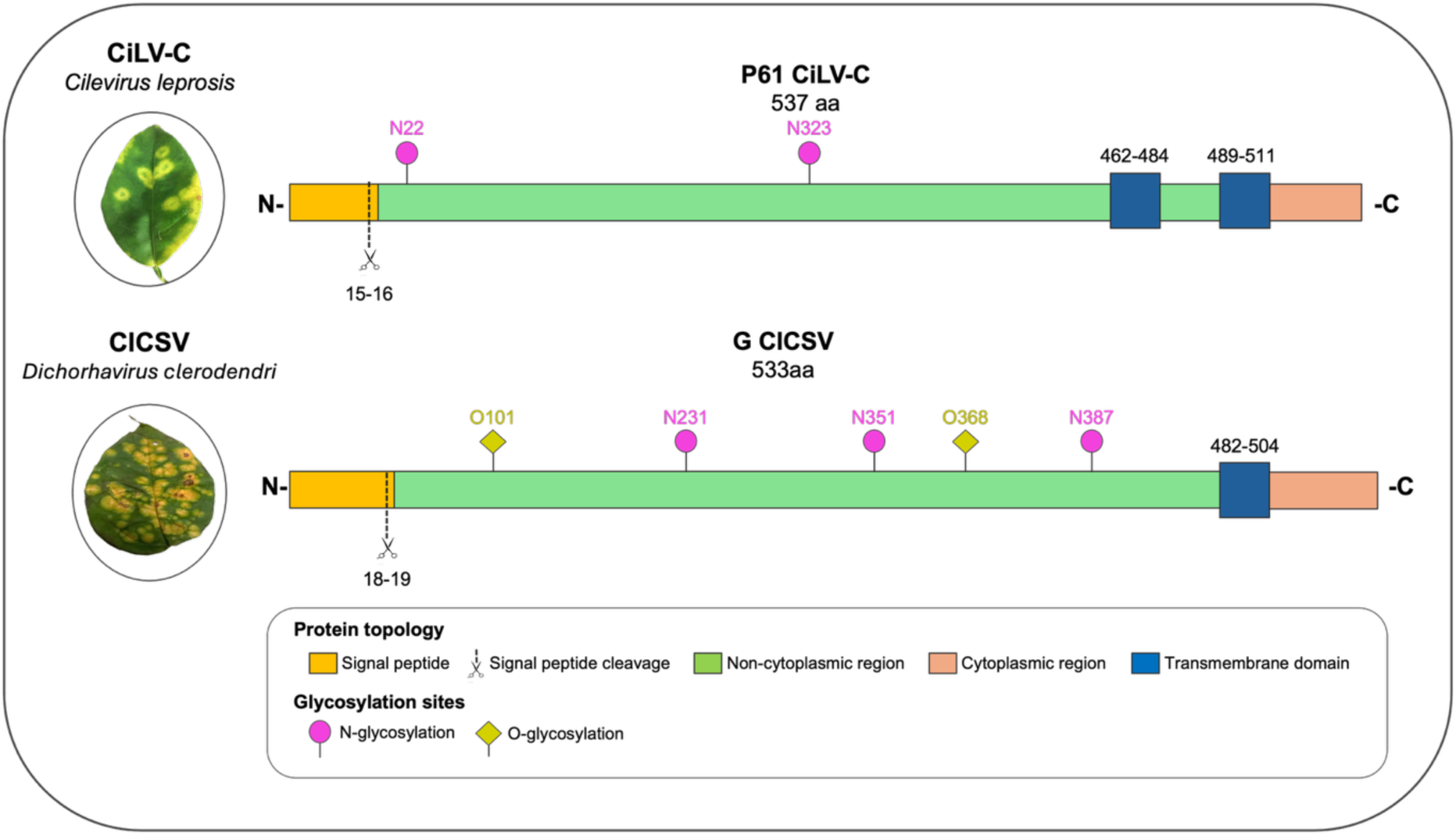
*In silico* characterization of the putative glycoproteins P61-CiLV-C and G-ClCSV. Representative sweet orange (*Citrus sinensis*) and *Clerodendrum* sp. leaf symptoms induced by CiLV-C (genus *Cilevirus*) and ClCSV (genus *Dichorhavirus*), respectively, are shown on the left. Linear diagrams depict the predicted structural organization of P61 and G, including the N-terminal signal peptide (yellow), predicted signal peptide cleavage site (scissors), non-cytoplasmic region (green), cytoplasmic region (salmon), transmembrane domain(s) (blue), predicted N-glycosylation sites (magenta circles), and predicted O-glycosylation sites (yellow diamonds). Signal peptides were predicted using SignalP 6.0 (SP probability: P61 = 0.99; G = 0.83), transmembrane topology using TMHMM 2.0 and InterPro, and N- and O-glycosylation sites using NetNGlyc 1.0 and NetOGlyc 4.0 (threshold = 0.5), respectively.

Together, these structural features, combined with the conserved genomic organization of cileviruses and dichorhaviruses, support the use of P61 and G as representative model glycoproteins for investigating the molecular basis of virus-vector interactions across both viral genera.

### MbY2H screens with P61 and G reveal numerous *B. yothersi* interacting proteins

To assess bait protein expression and establish optimal selection stringency for MbY2H screening, P61-CiLV-C and G-ClCSV were independently co-transformed with control prey plasmids in *Saccharomyces cerevisiae* NMY51. Growth occurred with the positive-control prey (pOst1-NubI), confirming correct bait expression, membrane targeting, and absence of autoactivation (Supplementary Figure S1). Under equivalent assay conditions, P61 consistently yielded fewer colonies than G across all 3-AT (3-amino-1,2,4-triazole, a competitive inhibitor of the *HIS3* reporter gene) concentrations tested, suggesting that P61 expression may impair yeast viability, consistent with P61-associated toxicity in *N*. *benthamiana* and Sf9 cells.

Library-scale screens using the *B. yothersi* cDNA library revealed striking differences in colony recovery between the two viral baits across tested stringencies. For P61, 1 mM 3-AT yielded 156 colonies, whereas 2.5 mM reduced recovery to only 20 colonies. In contrast, G remained highly productive even under higher stringency, generating 278 colonies at 5 mM and 1,000 colonies at 7.5 mM 3-AT. These results supported the use of low stringency (1 mM) for P61 and medium stringency (5 mM) for G to balance prey recovery with false-positive suppression. Together, these observations indicate that P61 imposes expression-associated constraints in yeast, likely reducing the recovery of low-abundance or transient interactors relative to G.

### Cilevirus and dichorhavirus glycoproteins exhibit distinct but partially overlapping interactomes in *B. yothersi*

MbY2H screening revealed asymmetrical interaction profiles for the two viral glycoproteins. G-ClCSV yielded nearly twice as many colonies as P61-CiLV-C, despite higher stringency, indicating an enhanced capacity of G to engage *Brevipalpus* proteins (Table 1). After redundancy removal, 162 unique interactors were identified for G and 73 for P61 (Table 1, Supplementary Data S1). Fewer than 5% of prey sequences corresponded to plant or microbial contaminants and these were excluded from subsequent analysis (Supplementary Table S2). Several additional sequences showed high identity to the *B. obovatus* genome (36) but were absent from the current *B. yothersi* assembly (34), likely reflecting incomplete genome annotation. These sequences were therefore retained as *Brevipalpus*-derived interactors and included in downstream analyses (Supplementary Table S2). The remaining interactors comprised 117 annotated, 12 hypothetical, and 5 uncharacterized proteins for G, and 37 annotated, 5 hypothetical, and 3 uncharacterized proteins for P61 (Figure 2). Notably, 13 annotated proteins, four hypothetical proteins, and one uncharacterized protein were shared between both viral glycoproteins, suggesting the existence of a conserved host interaction module.

**Figure 2.**
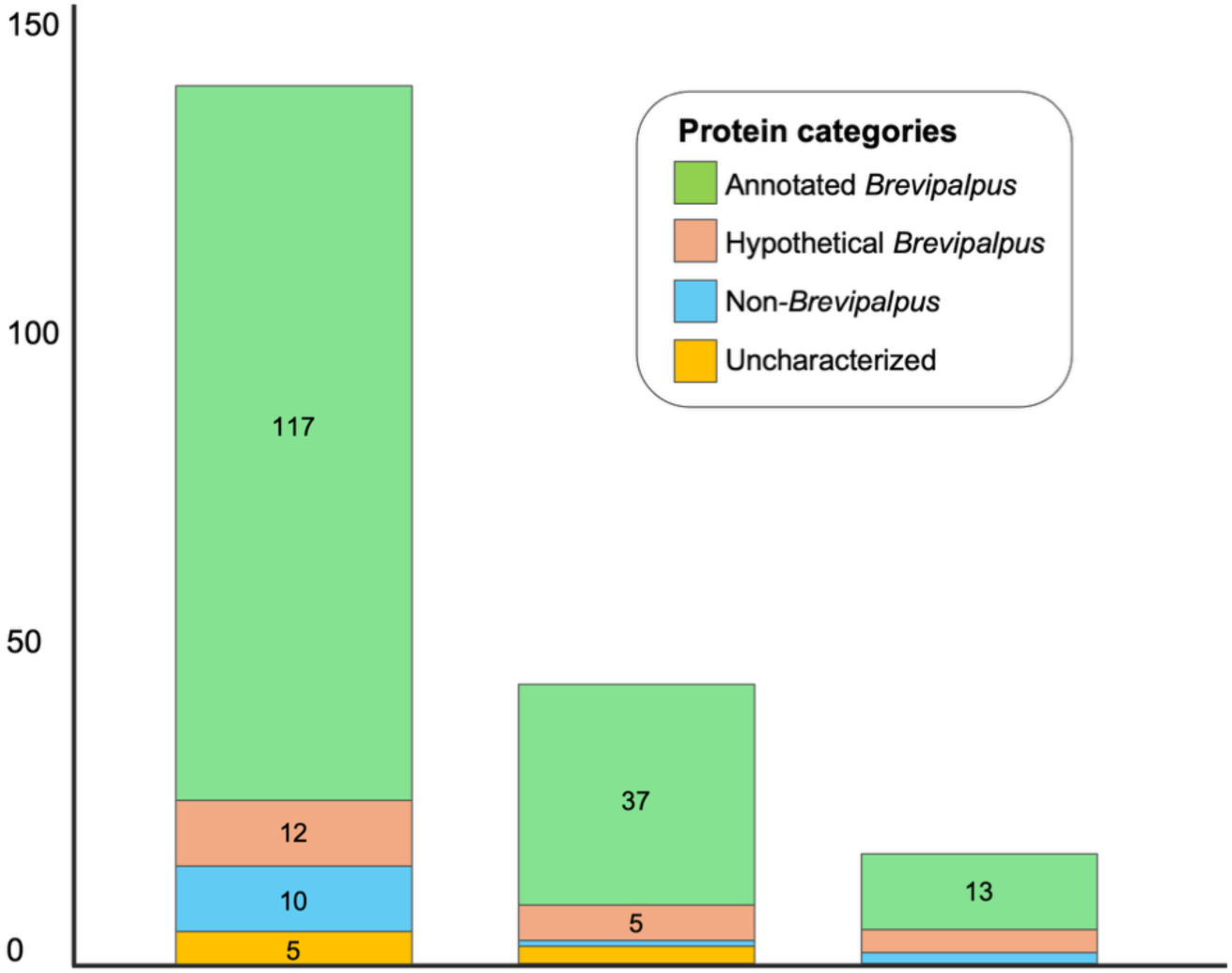
Classification of prey sequences based on *Brevipalpus* spp. genome annotations. Prey sequences matching *B. yothersi* or related *Brevipalpus* proteins (e.g., *B. obovatus*) were categorized as annotated proteins (green hypothetical proteins (salmon), or uncharacterized proteins (pink). Sequences with no detectable homology to *Brevipalpus* genomes, either matching other organisms or lacking clear taxonomic assignment, were classified as non-*Brevipalpus* sequences (blue). The chart illustrates the distribution of these categories for each bait interactor (P61 or G) and highlights the subset of prey proteins identified in both screens (Shared). Only categories containing ≥5 sequences are labeled in the graphs for clarity. Categories with <5 sequences were as follows: P61, 1 non-*Brevipalpus* and 3 uncharacterized proteins; Shared, 4 hypothetical proteins and 1 uncharacterized protein.

**Table 1.** Prey proteins identified in the MbY2H screens with P61-CiLV-C and G-ClCSV.

| Bait viral glycoprotein | Prey *Brevipalpus yothersi* proteins |  |  |  |
| --- | --- | --- | --- | --- |
|  | Positive colonies | Sequenced clones<sup>1</sup> | Non-redundant proteins | No significant hits |
| P61-CiLV-C | 158 | 115 | 73 | 6 |
| G-CiCSV | 286 | 247 | 162 | 20 |

<sup>1</sup>To maximize recovery of full-length coding sequences, only prey plasmids containing inserts larger than 0.9 kb were selected for Sanger sequencing.

Forty-three P61 and seventy-two G interactors were selected for validation based on membrane association, annotation quality, recurrence across clones, and prior evidence of viral interactions. Individual MbY2H assays confirmed growth on selective media for all tested candidates, although several interactions weakened under elevated stringency (Supplementary Table S2), consistent with transient or low-affinity binding.

Overall, the asymmetrical interactomes, broader for G and more restricted for P61, suggest distinct host engagement strategies by cileviruses and dichorhaviruses. These differences may reflect fundamental aspects of their infection biology within *B. yothersi*, whereas the shared subset of proteins likely represents a conserved host interaction module targeted by both viruses.

### P61 and G engage partially overlapping functional and subcellular classes of membrane-associated *B. yothersi* proteins

Functional categorization of annotated interactors using annotated proteins revealed broad enrichment in intracellular trafficking, secretion, and vesicular transport (COG U), as well as translation and ribosomal function (COG J) (Figure 3). G-ClCSV uniquely recovered proteins involved in chromatin organization (COG B), host defense (COG V), and cell-cycle control (COG D), whereas P61-CiLV-C recovered proteins related to secondary metabolite processes (COG Q) and DNA repair (COG L). Interactors common to both glycoproteins were predominantly associated with COG U and J, suggesting that trafficking and translational pathways constitute a shared functional core.

**Figure 3.**
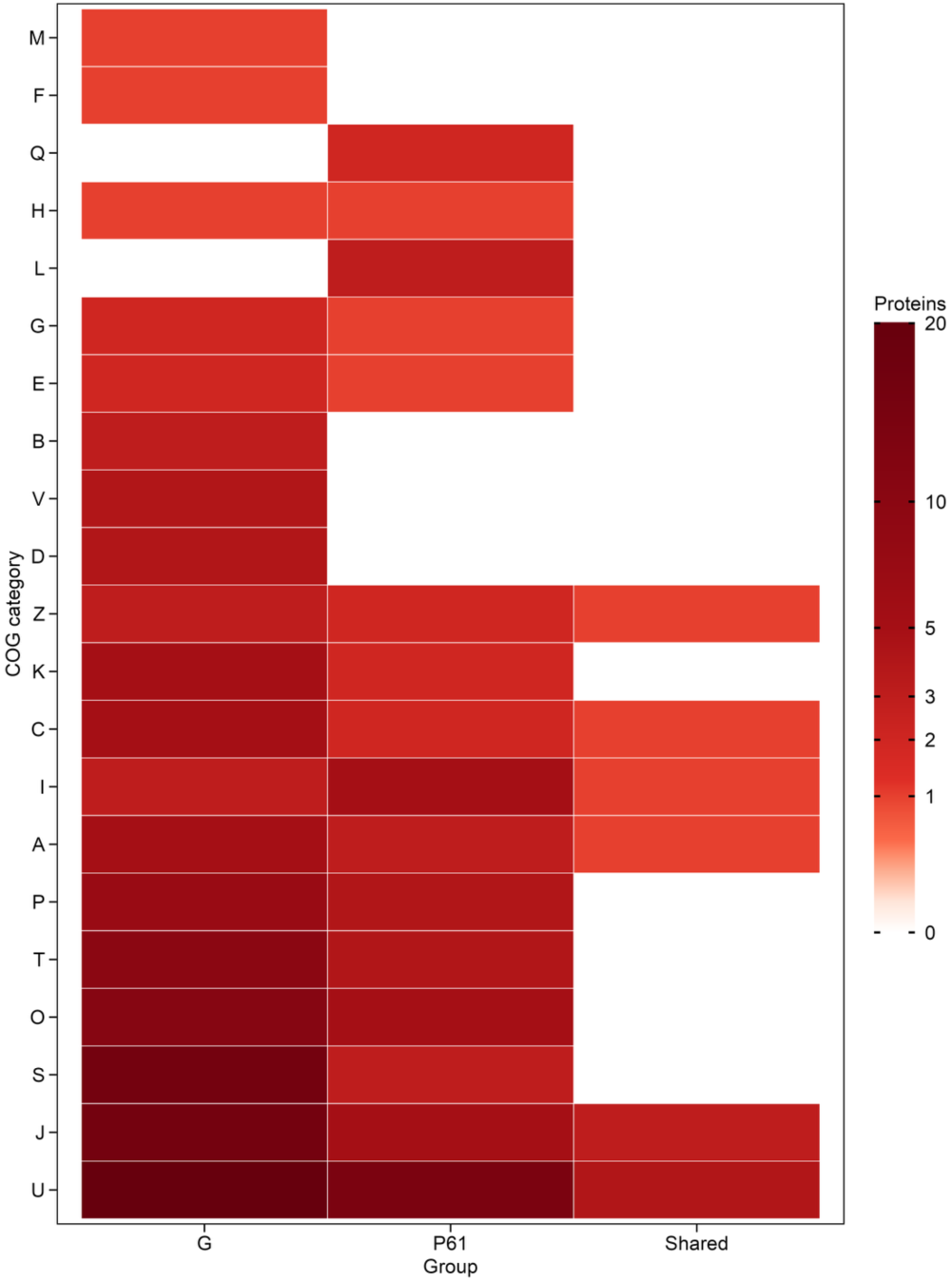
Heatmap of annotated *Brevipalpus* proteins interacting with G-ClCSV (G), P61-CiLV-C (P61), or both viral glycoproteins (both), categorized according to the database of Clusters of Orthologous Groups of proteins (COG). COG functional categories are defined as follows: U, Intracellular trafficking, secretion, and vesicular transport; J, translation, including ribosome structure and biogenesis; S, function unknown; O, molecular chaperones and related functions; T, signal transduction; P, inorganic ion transport and metabolism; A, RNA processing and modification; I, lipid metabolism; C, energy production and conversion; K, transcription; Z, cytoskeleton; D, cell division and chromosome partitioning; V, defense mechanisms; B, chromatin structure and dynamics; E, amino acid transport and metabolism; G, carbohydrate transport and metabolism; L, replication, recombination, and repair; H, coenzyme metabolism; Q, secondary metabolites biosynthesis, transport and catabolism; F, nucleotide transport and metabolism; M, cell wall structure, biogenesis, and outer membrane;

DeepLoc-based subcellular predictions showed that ∼84–85% of interactors were membrane-associated, with strong ER and nuclear envelope enrichment (Figures 4–5). G-specific interactors showed higher representation of nuclear envelope proteins, whereas P61 interacted with a more even distribution across ER, plasma membrane, Golgi, and mitochondria. Proteins shared by both viral glycoproteins were enriched in ER–Golgi trafficking components.

**Figure 4.**
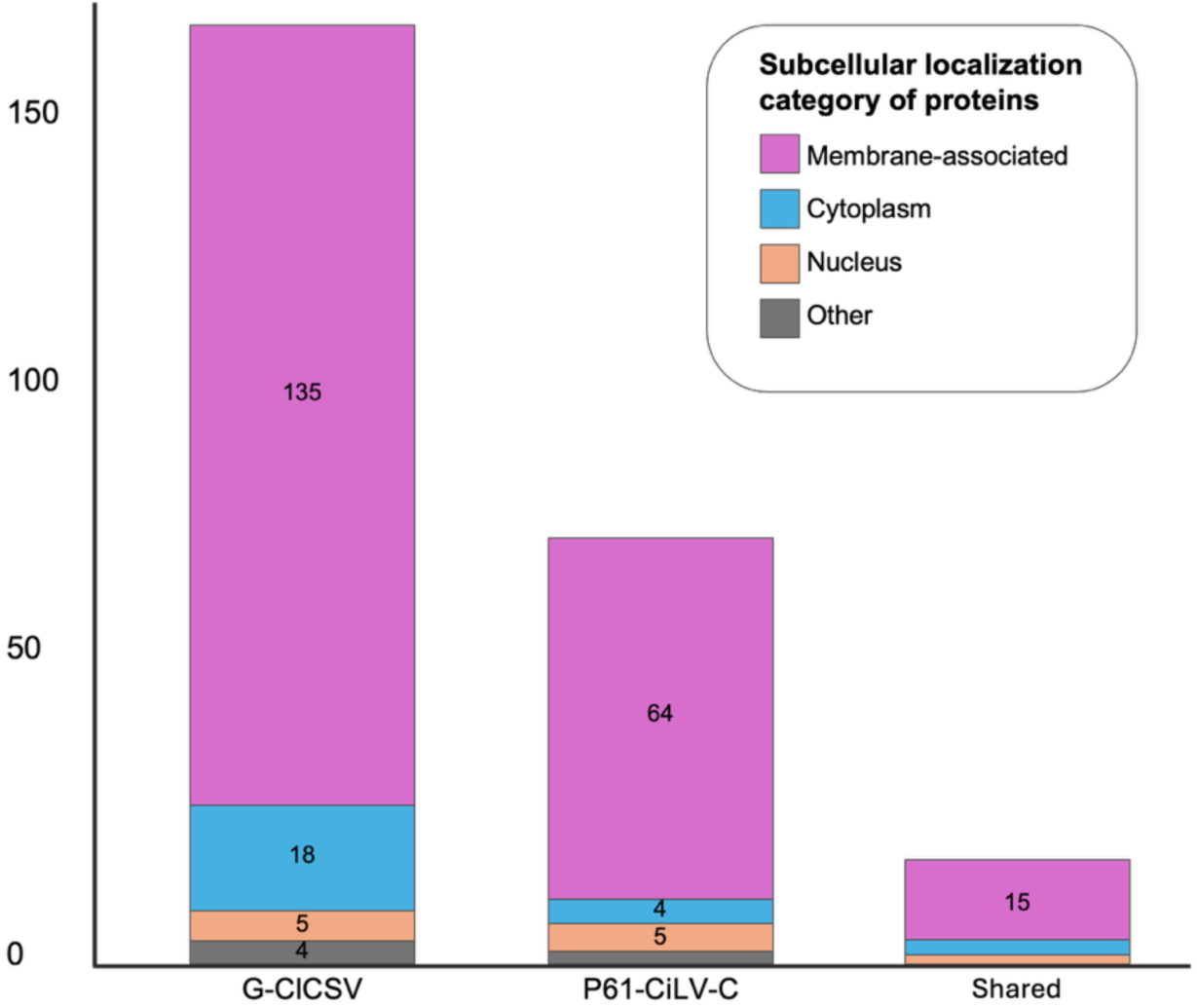
Predicted subcellular localization of annotated *Brevipalpus* proteins interacting with G-ClCSV (G), P61-CiLV-C (P61), or both viral glycoproteins (both), based on DeepLoc 2.0 prediction. Annotated proteins were classified as transmembrane or membrane-associated (pink), cytoplasmic (blue), nuclear (salmon), or localized to other cellular compartments (grey). Only categories containing ≥4 sequences are labeled in the graphs for clarity. Categories with <4 sequences were as follows: P61, 2 proteins classified as Other; Shared, 2 cytoplasmic proteins and 1 nuclear protein.

**Figure 5.**
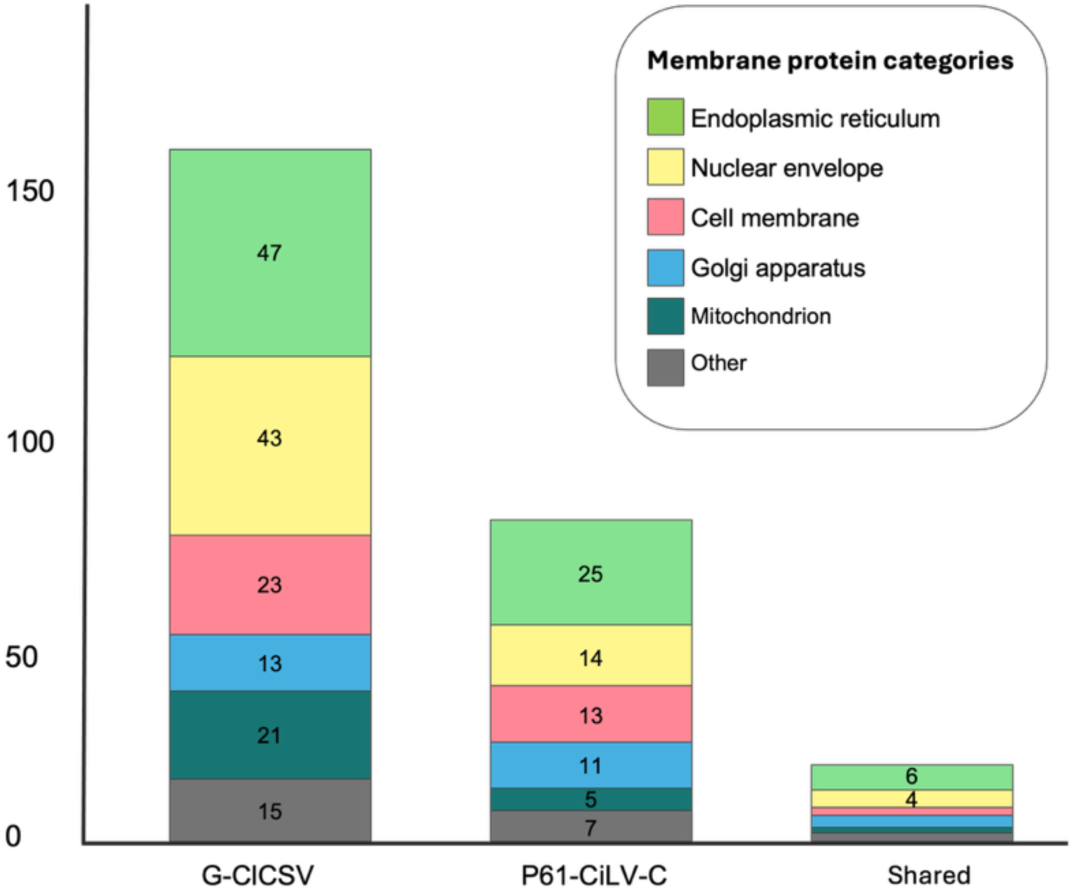
Predicted subcellular membrane localization of transmembrane or membrane-associated *Brevipalpus* proteins interacting with viral glycoproteins. Subcellular localization prediction using DeepLoc 2.0 revealed the distribution of these proteins across membrane-associated compartments, including the endoplasmic reticulum (green), nuclear envelope (yellow), cell membrane (salmon), golgi apparatus (blue), mitochondrion (teal) and other locations (grey). Only categories containing ≥4 sequences are labeled in the graphs for clarity. Categories with <4 sequences were as follows: Shared, 2 cell membrane, 3 Golgi apparatus, 1 mitochondrion and 2 other location.

Hypothetical and uncharacterized proteins (n=29 total) were overwhelmingly predicted to encode signal peptides and/or transmembrane domains (∼90%) (Figure 6), supporting potential roles as receptor-like or membrane-anchored determinants of virus–vector specificity. InterProScan identified an ApoLp-III domain in bryot108g00160 (G-specific) and coiled-coil motifs in some G and P61 interactors.

**Figure 6.**
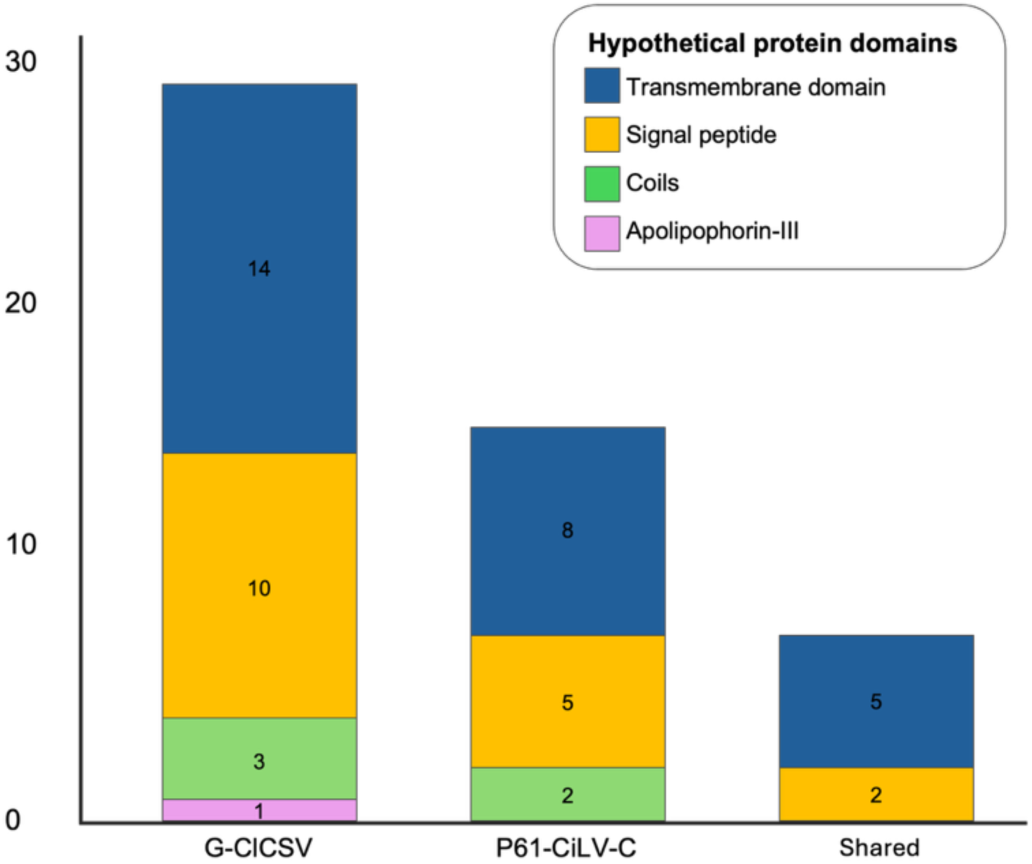
Predicted protein domains in hypothetical or uncharacterized *Brevipalpus* proteins interacting with viral glycoproteins identified in the MbY2H screen. Domain prediction using InterProScan revealed the presence of signal peptides (red), transmembrane domains (blue), Apolipophorin-III–like domains (green) and coiled-coil regions (coils, purple) among the *Brevipalpus* proteins that interact with viral glycoproteins (G, P61 or both).

Together, these findings indicate that both glycoproteins interact with proteins associated with ER- and nuclear envelope localized pathways, with additional Golgi-associated factors reflecting broader engagement of the early secretory system. G displayed a proportionally stronger association with nuclear envelope proteins, whereas P61 showed a more evenly distributed membrane profile.

### Structural modeling predicts well-defined folds in *B. yothersi* proteins and lower-confidence structures for viral glycoproteins

Structural modeling was performed to examine the three *B. yothersi* proteins selected for individual validation experiments: ADP-ribosylation factor 1 (ARF1; 181 aa), stress-associated endoplasmic reticulum protein 2 (SERP2; 64 aa), and the hypothetical transmembrane protein bryot216g00140 (156 aa). These proteins represented robust and recurrent interactors in the MbY2H screen and were recovered multiple times with both viral glycoproteins (Supplementary Table S3).

AlphaFold modeling yielded folded α-helical architectures for all three proteins, with high-confidence predictions across most residues (pLDDT 70–90) (Figure 7). SERP2 and bryot216g00140 each contained predicted transmembrane helices with strong N-in orientation, and bryot216g00140 displayed a coiled-coil–like configuration of helices.

**Figure 7.**
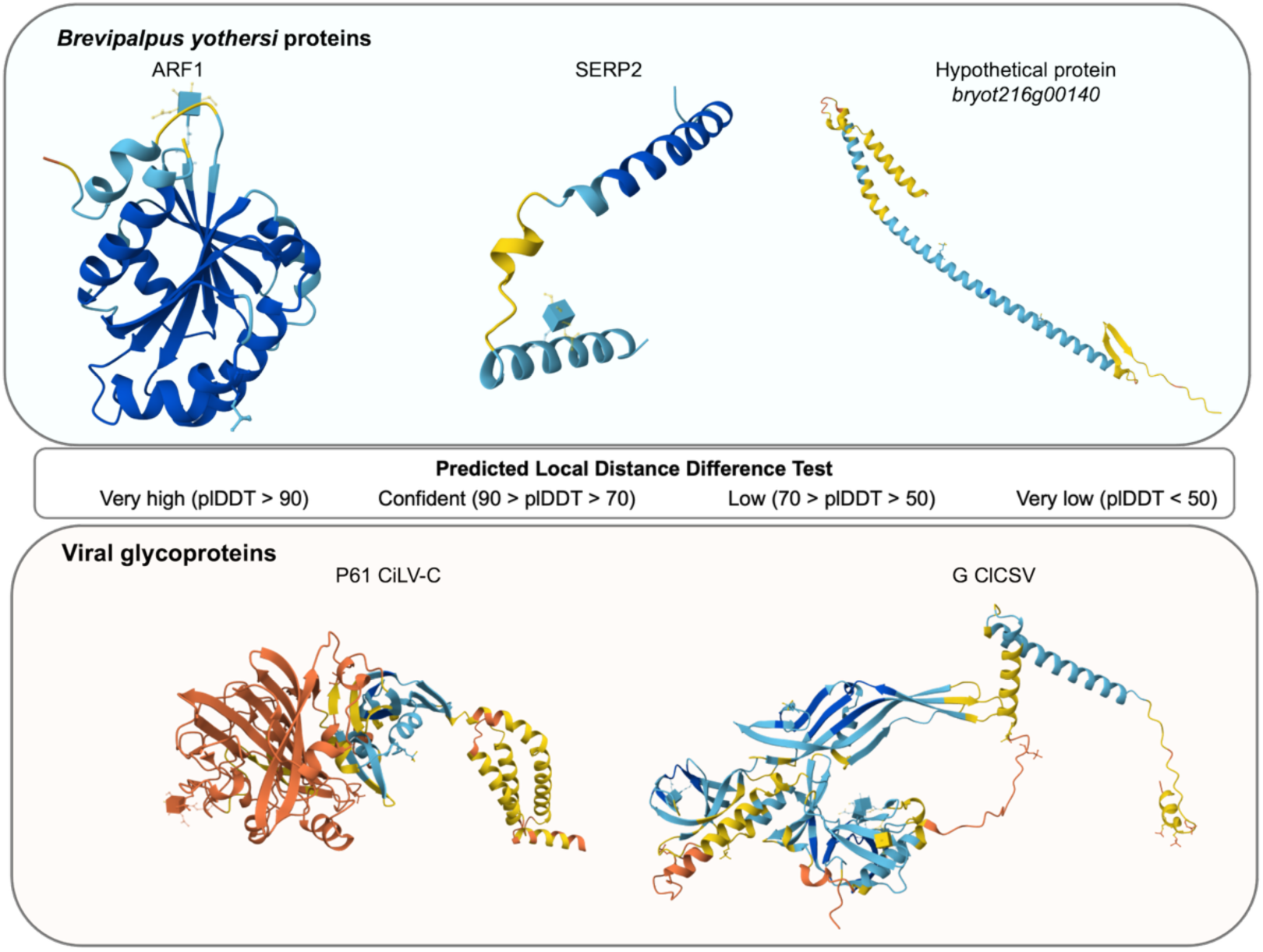
Three-dimensional structural predictions of *Brevipalpus yothersi* proteins and the viral glycoproteins P61 and G generated with AlphaFold 3.0. Signal peptide regions were removed prior to modeling, and relevant post-translational modifications were incorporated into the final predicted structures. Predicted Local Distance Difference Test (pLDDT) is a per-residue confidence score used by AlphaFold to estimate the reliability of structural predictions.

In contrast, the viral glycoproteins G-ClCSV and P61-CiLV-C showed mixed α-helical, β-strand, and extended coil regions interspersed with areas of predicted disorder and lower-confidence modeling (Figure 7). Such patterns are commonly observed for viral and transmembrane proteins in AlphaFold predictions due to the limited availability of close structural homologs (47), and therefore cannot be interpreted as evidence of functional dynamics or conformational flexibility. Likewise, protein–protein complex predictions did not yield high-confidence viral–host interaction models (ipTM/pTM < 0.4) (Supplementary Fig. S2; Supplementary Tables S3), likely reflecting these same limitations in structural information rather than the absence of interaction potential.

Together, these analyses indicate that the *B. yothersi* proteins possess defined structural folds consistent with membrane-associated roles, whereas the lower-confidence and partially disordered regions predicted for the viral glycoproteins most likely reflect constraints inherent to structural prediction for viral and transmembrane proteins.

### Sf9 expression assays confirm membrane association and differential stability of viral and *B. yothersi* mite proteins

Transient expression of GFP-tagged viral proteins in Sf9 cells showed cytoplasmic and perinuclear localization consistent with ER–Golgi association (Figure 8). G-ClCSV was readily detectable from 24–120 hours post-transfection (hpt), whereas P61 fluorescence was transient and limited to a few cells, a pattern that may reflect progressive cytotoxicity (Supplementary Figure 3). Signal peptide and transmembrane deletion mutants failed to accumulate detectable fluorescence (except for a very weak signal in the G-ΔTM mutant), demonstrating that these domains are essential for stability and expression in insect cells (Supplementary Figure 4).

**Figure 8.**
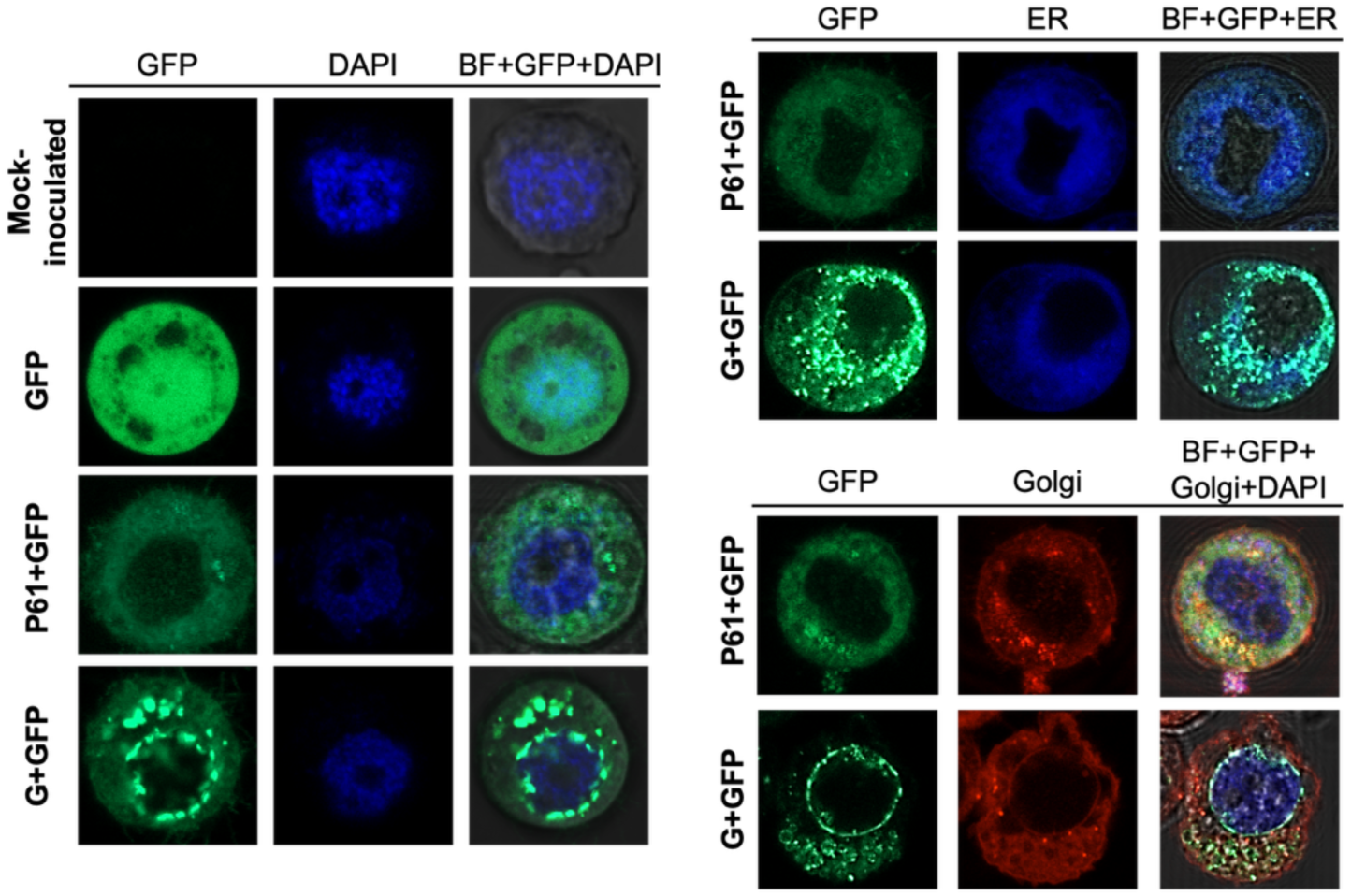
Heterologous transient expression of P61-CiLV-C and G-ClCSV proteins in Sf9 cells. The viral glycoproteins were fused to GFP, and fluorescence images were acquired at 24 hours post-transfection for P61 and 72 hours post-transfection for G using a Zeiss LSM 980 confocal microscope. Sf9 cells without plasmid (mock-inoculated) were used as the negative control, and cells transfected with the empty pI-GFP vector expressing free GFP served as the positive control (GFP). To assess subcellular localization, nuclei were labeled with DAPI (blue) and established markers for the endoplasmic reticulum (ER, blue) and Golgi apparatus (Golgi, red) were included. Merged images incorporate the brightfield (BF) channel to visualize cellular morphology.

RFP-tagged SERP2 and ARF1, and GFP-tagged bryot216g00140, localized to cytoplasmic and ER/Golgi-associated regions (Figure 9). SERP2 overlapped strongly with ER markers, ARF1 localized to the cytoplasm and nucleus, and bryot216g00140 accumulated in perinuclear regions.

**Figure 9.**
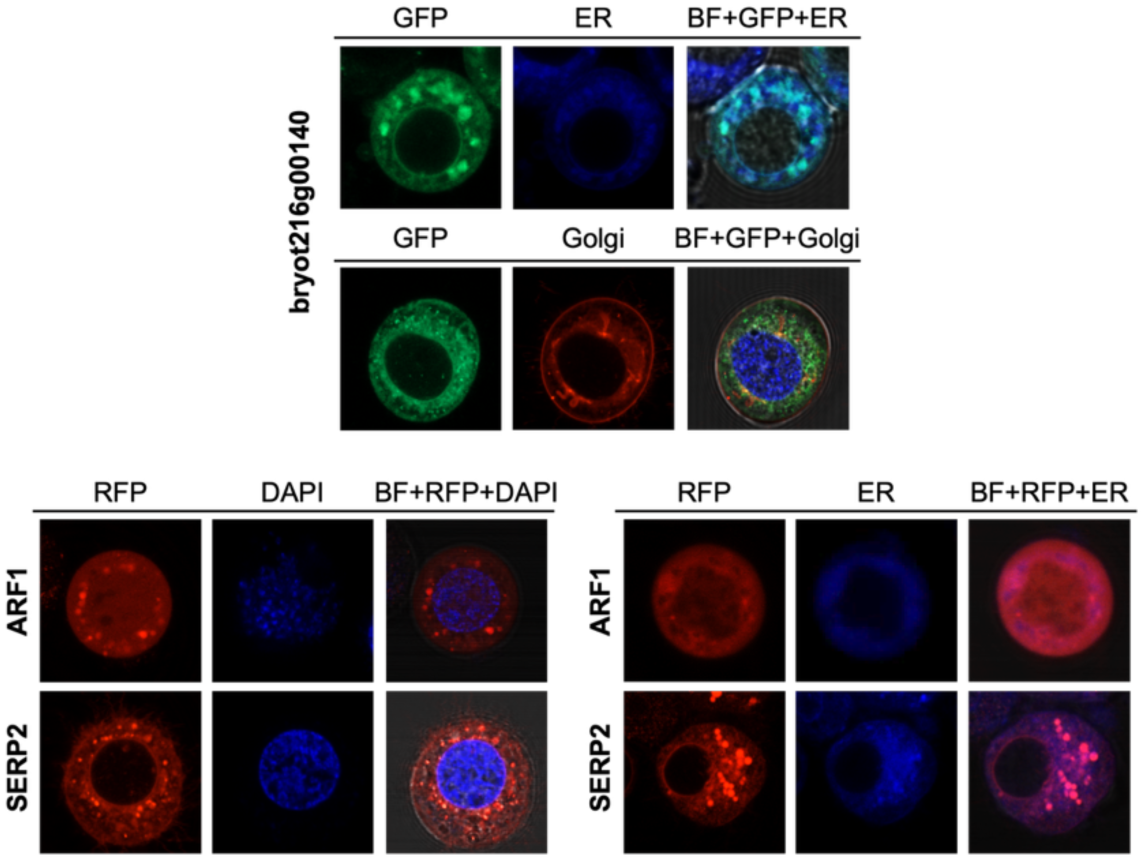
Heterologous transient expression of *Brevipalpus yothersi* proteins in Sf9 cells. The hypothetical *B. yothersi* protein (bryot216g00140) was fused to GFP, whereas the annotated proteins ARF1 and SERP2 were fused to RFP. Fluorescence images were acquired 72 hours post-transfection using a Zeiss LSM 980 confocal microscope. When applicable, nuclei were stained with DAPI (blue), and established markers for the endoplasmic reticulum (ER, blue) and Golgi apparatus (red) were included. Merged images incorporate the brightfield (BF) channel to visualize overall cell morphology.

Thus, Sf9 expression assays demonstrate conserved membrane targeting across viral and mite proteins while revealing intrinsic instability of P61, underscoring functional differences between the two glycoproteins.

### BiFC and Co-IP confirm physical interactions between BTV glycoproteins and *B. yothersi* proteins in arthropod cells

BiFC assays demonstrated strong YFP reconstitution for G-ClCSV co-expressed with ARF1, SERP2, or bryot216g00140 in Sf9 cells (Figure 10). The BiFC signal accumulated in regions where the host proteins are typically enriched, including perinuclear domains for bryot216g00140 and ER-like reticulation for SERP2. ARF1-associated BiFC fluorescence remained predominantly cytoplasmic when co-expressed with G, indicating that the interaction occurs mainly within this compartment. For P61-CiLV-C, YFP signal was detected only with bryot216g00140, appearing transiently at ∼22 hpt (Figure 10), consistent with P61-induced cytotoxicity and limited protein accumulation.

**Figure 10.**
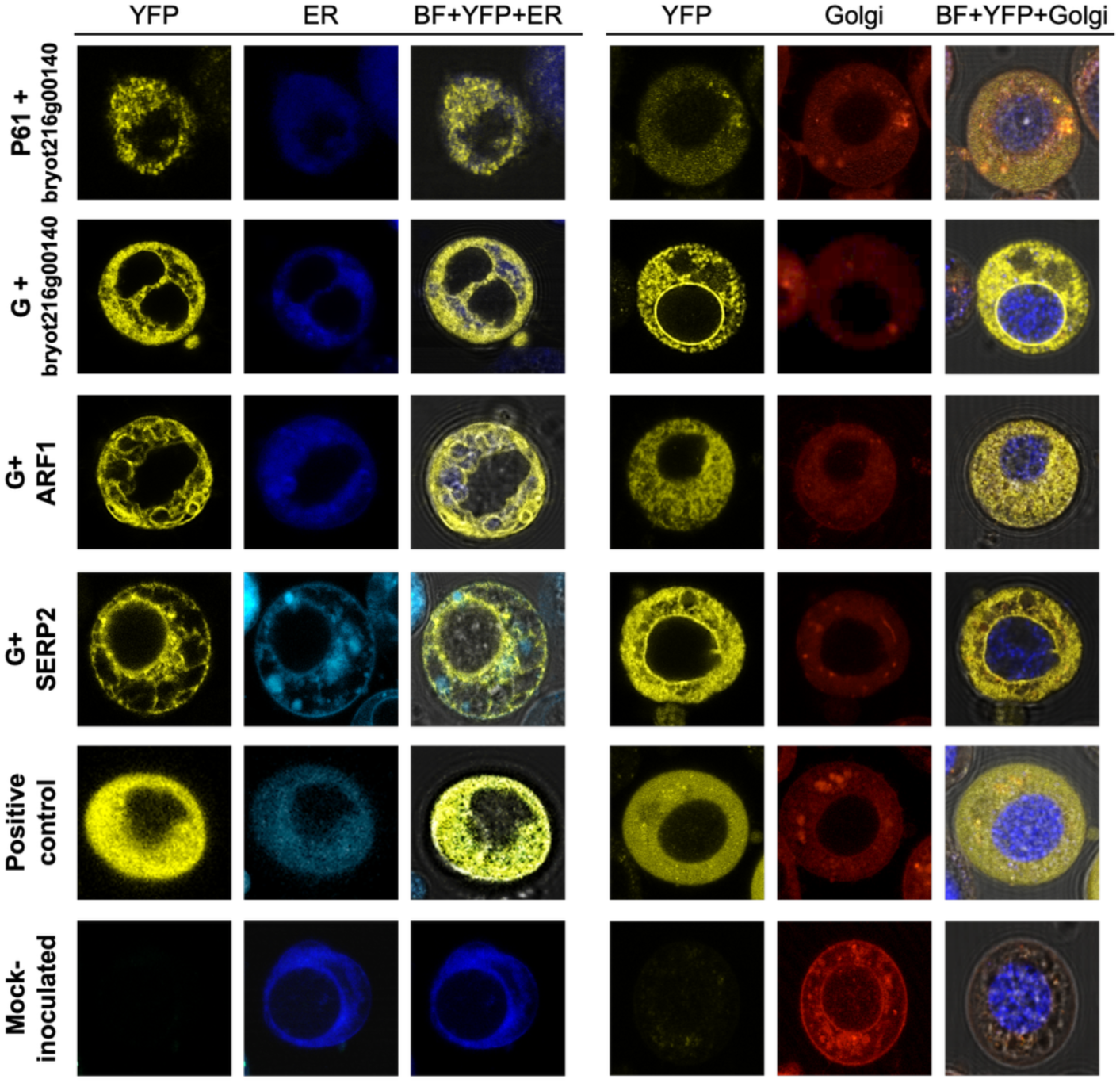
Bimolecular fluorescence complementation assay of interactions between *B. yothersi* proteins and BTV glycoproteins in Sf9 cells. Sf9 insect cells were transfected with dual-promoter BiFC constructs expressing the viral glycoproteins fused to cYFP and mite candidate proteins fused to nYFP. Cells were examined by confocal microscopy (Zeiss LSM 980) at 24 h post-transfection for P61 interactions and at 72 h for G interactions. Reconstituted YFP fluorescence indicates physical association between the mite protein and either P61 or G. Subcellular localization was assessed using DAPI for nuclei (blue), an endoplasmic reticulum marker (ER; blue), and a Golgi apparatus marker (Golgi; red). Merged panels include the brightfield (BF) channel to visualize cell architecture. Negative controls consisted of non-transfected Sf9 cells, whereas positive controls included cells expressing cYFP–MBP and nYFP–MBP, a previously characterized self-interacting protein.

Co-IP assays confirmed all interactions detected by BiFC, including P61 interactions with SERP2 and ARF1 (Figure 11). Using anti-FLAG enrichment of viral proteins and anti-Myc detection of mite proteins, ARF1, SERP2, and bryot216g00140 co-precipitated specifically with their respective viral partners.

**Figure 11.**
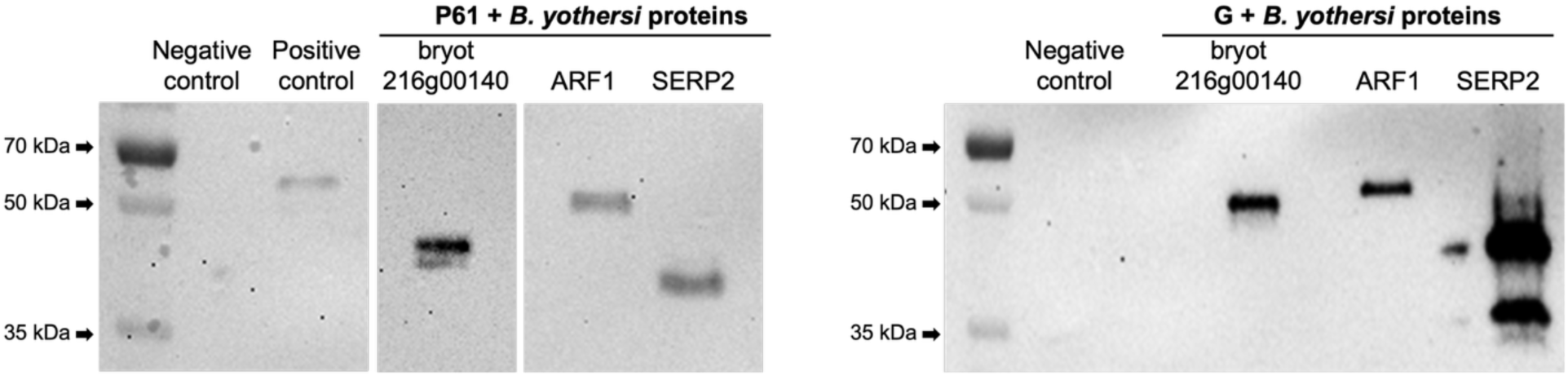
Co-immunoprecipitation analysis of *B. yothersi* proteins fused to Myc and their association with BTV glycoproteins in Sf9 cells. Sf9 cells were harvested at 24 h post-transfection for P61 assays and at 72 h for G assays. Approximately 2 × 10⁶ (P61 interactions) or 1 × 10⁶ (G interactions) transfected Sf9 cells were collected for protein extraction and immunoprecipitation using Anti-FLAG Affinity Gel. Western blot detection of co-immunoprecipitated mite proteins was performed using an anti–c-Myc monoclonal antibody (Thermo Fisher Scientific), followed by Mouse TrueBlot® ULTRA Anti-Mouse Ig HRP (Rockland) as the secondary antibody. The expected molecular masses of the mite protein–Myc fusions were 47 kDa for the hypothetical protein bryot216g00140, 50 kDa for ARF1, and 37 kDa for SERP2. Negative controls consisted of co-expression of the viral glycoprotein–cYFP vector (P61 or G, depending on the experiment) with nYFP–MBP, which does not interact with the viral proteins. Positive controls included co-expression of cYFP–MBP and nYFP–MBP, a known self-interacting protein, which generates a ∼70 kDa band corresponding to the MBP–nYFP fusion. Protein ladder: Thermo Scientific™ PageRuler™ Plus Prestained Protein Ladder, 10 to 250 kDa (Thermo Fisher Scientific).

Together, the BiFC and Co-IP assays validate ARF1, SERP2, and bryot216g00140 as bona fide interactors of BTV glycoproteins in arthropod cells, providing strong experimental support for the interactome derived from MbY2H screening.

## DISCUSSION

Enveloped plant viruses are exceedingly rare, and their membranes contribute little to infection in walled plant cells (48,49). Instead, increasing phylogenetic, ultrastructural, and cellular evidence indicates that these membranes reflect deep and persistent associations with arthropod hosts, where replication, morphogenesis, and host-virus interactions occur far more extensively than in plants (48). *Brevipalpus*-transmitted viruses (BTVs) exemplify this biology: although enveloped, they induce strictly localized, non-systemic lesions in plants, yet replicate, assemble, and persist across multiple mite tissues and developmental stages (8). Emerging observations that BTV infection may even enhance mite performance further reinforce the view that *Brevipalpus* mites represent the primary adaptive environment for these viruses, whereas plants serve mainly as temporary transmission platforms (8,24). Within this framework, understanding the molecular role of viral glycoproteins, long suspected to function predominantly in vector tissues, becomes essential for elucidating transmission. Here, we test a key prediction of this vector-centric model, namely that the putative glycoproteins of cileviruses (P61) and dichorhaviruses (G) have evolved to interact with mite cellular machinery, particularly pathways involved in intracellular trafficking, membrane remodeling, and receptor recognition.

Among the structural proteins putatively associated with the viral envelope, the cilevirus P61 has received the most attention (8) *In planta*, P61 localizes to the ER, disrupts secretory pathway architecture, induces ROS accumulation, perturbs SA/JA signaling, and triggers HR-like cell death reminiscent of citrus leprosis lesions (25,50). In contrast, far less is known about the specific glycoprotein of dichorhaviruses, although extensive studies of plant and animal rhabdovirids provide important context (7,21,27). Phylogenetic clustering of dichorhavirus G proteins according to *Brevipalpus* species groups (29), along with their nuclear envelope localization *in planta* (51), provides indirect evidence supporting the hypothesis that G functions as the dichorhavirus glycoprotein. Functional parallels with rhabdoviridal glycoproteins, including roles in trafficking, nuclear envelope remodeling, and autophagy-associated responses (21,28,52), further support the idea that G contributes centrally to vector specificity. Taken together, if these proteins indeed function as the structural glycoproteins of their respective viruses, both P61 and G could participate in processes relevant to interactions within the mite vector.

A major conceptual shift introduced by this work is the transition from a plant-centered to a vector-centered view of BTV glycoprotein function. According to the vector-centric model, these glycoproteins are expected to interact with mite proteins involved in pathways that viruses typically exploit during entry, intracellular movement, assembly, and egress, such as vesicular trafficking, ER-Golgi networks, and receptor-mediated recognition (21,53). By integrating MbY2H screening with *in vivo* validation in Sf9 insect cells, we map, for the first time, the interactomes of P61 and G within an arthropod-like intracellular environment. Although heterologous, Sf9 cells preserve key secretory pathway features of arthropods, enabling detection of interactions that may be absent, masked, or irrelevant in plants. This approach aligns with recent vector-focused interactome studies conducted for other plant rhabdovirids, such as maize mosaic virus (MMV, *Alphanucleorhabdovirus maydis*), in which MbY2H screening revealed vector proteins associated with endocytosis, trafficking, metabolism, and defense (54,55). Our results extend this paradigm to BTVs and uncover a dual pattern of divergence and convergence: P61 and G exhibit distinct interaction profiles yet share a conserved subset of *Brevipalpus* proteins that cluster within similar functional categories. This shared core strongly suggests convergent adaptation driven by the common vector environment (6).

When expressed in Sf9 cells, both P61 and G displayed diffuse cytoplasmic localization with perinuclear accumulations consistent with ER–Golgi association (Figure 8). This distribution aligns with the expected maturation route of viral glycoproteins and with the enrichment for ER/Golgi-associated factors recovered in the interactomes (27,56). P61 expression, however, produced rapid cell death in both yeast and insect cells, mirroring its documented cytotoxicity in plants (25,50). Although the mechanisms underlying toxicity in arthropods remain unknown, viral glycoproteins from arthropod-borne systems often induce ER stress, disrupt trafficking, activate UPR pathways, and cause misfolding-associated proteotoxicity (28,56,57). Whether P61 toxicity reflects native infection processes or overexpression artifacts remains to be determined, but its strong cytopathic signature underscores the need to explore whether CiLV-C modulates ER homeostasis in mites as it does in plants (25).

The interactome sizes differed substantially: 73 interactors for P61 and 162 for G (Table 1). This disparity likely reflects biological differences in the intracellular life cycles of cileviruses and dichorhaviruses within mites. CiLV-C has never been conclusively observed replicating within mite cells; virions are typically detected in paracellular spaces of podocephalic glands, ovaries, and muscles (8,16,58). Although antigenomic RNA persists in mites even when they feed on noninfected plants, viral replication appears limited, restricted to specific tissues or cell types (8). In contrast, ClCSV follows a clear propagative cycle characteristic of other dichorhaviruses(19): virions and viroplasms accumulate throughout multiple tissues (20,22,59), viral titers increase across developmental stages, and transmission requires long acquisition periods (A.D. Tassi, unpublished data). Such pervasive intracellular presence likely exposes G to a broader range of host machinery, contributing to the larger interactome. Additionally, P61-associated cytotoxicity in yeast (Supplementary Figure S1) may further reduce the recovery of P61 interactors. These differences are consistent with the vector-centric model: a virus that undergoes more extensive replication and intracellular trafficking within the mite (ClCSV) is expected to engage a broader range of host factors than one with a more restricted presence (CiLV-C). Thus, interactome size is consistent and provides indirect support for differential adaptation to the vector environment.

Functional classification revealed enrichment for intracellular trafficking, secretion, and vesicular transport (COG U) across both interactomes, and especially among the shared interactors (Figure 3). This finding directly supports the vector-centric model in which enveloped arthropod-borne viruses typically rely on host trafficking pathways for entry, intracellular movement, assembly, and egress (50,60). The convergence of both BTV glycoproteins on these same pathways suggests that, despite their distinct evolutionary histories, they have adapted to engage a conserved core of mite trafficking machinery. Additional enriched categories included translational machinery (J), post-translational modification and chaperones (O), signal transduction (T), and ion/lipid transport (I, P). These findings suggest that BTV glycoproteins rely on, and possibly modulate, components of protein synthesis, folding, and membrane remodeling within mite cells.

Among trafficking regulators (COG U), Rab GTPases were prominent. Rab14 interacted with both glycoproteins; Rab18 with G-ClCSV; and Rab8A, Rab27A, and Rab escort protein 1 with P61 (Supplementary Table S2). Rabs are essential for vesicle identity, directed trafficking, ER–lipid droplet contacts, and secretory vesicle movement (61). Notably, Rab14 was also recovered as a G-interactor in MMV (54), suggesting conserved viral targeting of Rabs across unrelated arthropod-borne viruses. Likewise, clathrin light chain B (CLTB), a core component of clathrin-mediated endocytosis (62), interacted specifically with G-ClCSV. Many enveloped viruses, including rhabdovirids, exploit clathrin-associated pathways for entry or intracellular movement (60,62–64), raising the possibility that dichorhaviruses engage similar pathways during acquisition or replication in mites. The repeated recovery of these trafficking regulators, which are conserved across eukaryotes (61), reinforces the idea that BTV glycoproteins have evolved to interface with ancient, arthropod-conserved pathways. Considering the vector-centric view, these interactions are unlikely to be incidental, and instead they likely contribute to viral recognition, internalization, and intracellular movement within the mite.Ribosomal proteins (RPs) represented ∼12% of annotated interactors (COG J), including both large and small subunits. RPs are increasingly recognized as modulators of virus–host interactions in plants and arthropods (65–69). Although their role in *Brevipalpus* remains unknown, their repeated recovery here suggests that BTV glycoproteins may interface with the translational machinery, possibly influencing viral protein synthesis or stress-associated remodeling of ribosomal function.

Category O proteins, including those involved in posttranslational modification and chaperone systems, were also well represented (10%) (Figure 3, Supplementary Table S2). Two glycosylation-associated proteins, RPN2 and NGC, interact specifically with G. RPN2 participates in early steps of N-glycosylation, whereas NGC is associated with chondroitin sulfate formation and has been implicated in viral attachment (70,71). On the other hand, the mitochondrial chaperone DNAJA3-like protein interacted specifically with P61. DnaJ/Hsp40 proteins regulate Hsp70 activity and can play antiviral or proviral roles across diverse systems (72,72–76). Their presence here suggests that both glycoproteins may modulate protein folding and quality control pathways during infection.

One notable outcome in this study was the recovery of numerous hypothetical and uncharacterized *Brevipalpus* proteins (21 hypothetical and 8 uncharacterized), most bearing signal peptides and/or transmembrane domains (Figures 2 and 6). Such features are typical of receptor-like proteins (77), making them strong candidates for mediating virus–vector specificity. An ApoLp-III domain was also predicted in the G-specific interactor bryot108g00160 (Supplementary Table S2), consistent with the known interaction between vector ApoLp-III and the G-MMV (54). Of particular interest is bryot216g00140, repeatedly recovered with both glycoproteins and validated by BiFC and Co-IP (Figures 10 and 11). Predicted as a two-pass membrane protein with N-in topology, it localizes to perinuclear/ER regions and may participate in glycoprotein recognition or trafficking. Its robust interaction with P61, despite P61-induced cytotoxicity, suggests a stable or high-affinity association. Collectively, these hypothetical and uncharacterized proteins, many bearing features of receptors or membrane-associated factors, represent strong candidates for mediating virus-vector specificity. Their identification provides a molecular starting point for understanding how BTVs recognize and enter mite cells.

The interaction of BTV glycoproteins with two annotated proteins (ARF1 and SERP2) was likewise validated. ARF1 is a master regulator of vesicle budding, Golgi function, and actin dynamics, supporting viral replication and spread in diverse arthropod and plant systems (78,79). SERP2 participates in ER-associated translocation and protects nascent proteins from misfolding (80), and related SERP family members modulate viral infection in insects (81). These interactions reinforce the central role of ER/Golgi pathways during BTV acquisition and intracellular trafficking, and further support the conclusion that the vector-centric model accurately predicts which host pathways are most relevant to BTV glycoprotein function.

Collectively, our findings support a vector-centric model in which BTV glycoproteins engage a network of *Brevipalpus* cell factors involved in vesicular trafficking, ER/Golgi processes, chaperone regulation, translation, and receptor-like membrane proteins. These interactions may facilitate viral recognition, stabilization, intracellular movement, and possibly assembly within vector tissues. The broader interactome of G-ClCSV is consistent with its propagative replication in mites, whereas the more restricted P61-CiLV-C interactome may reflect limited intracellular presence and/or the cytotoxic effects associated with P61 overexpression in heterologous systems. In contrast, plant hosts, where infection is localized, non-systemic, and non-essential for viral persistence, appear to play a secondary role, serving mainly as a transmission platform rather than a primary site of viral adaptation (17,82).

Although this study has limitations, including the use of yeast and insect cells that cannot fully reproduce *Brevipalpus* biology, the interactome generated here provides a framework for interpreting how BTV glycoproteins interface with components of the vector’s intracellular environment. The convergence of interactors between the two viral families suggests shared pathways shaping virus adaptation to *Brevipalpus* mites, whereas divergence highlights virus-specific strategies. Importantly, P61 has not yet been biochemically confirmed as a glycoprotein, and thus its functional classification remains inferred. Nevertheless, the interaction patterns observed here, particularly with membrane-associated and trafficking-related host factors, are consistent with a glycoprotein-like role and provide indirect support for its proposed biochemical nature.

Future experiments targeting key interactors through gene silencing or CRISPR-based approaches in mites will be essential to clarify their functional relevance during viral acquisition, replication, and transmission. Moreover, the convergence of our findings on pathways predicted by the vector-centric model provides a strong rationale for prioritizing these candidates in future functional studies directly in mites.

Taken together, these findings reveal that the likely structural glycoproteins of cileviruses and dichorhaviruses engage conserved host processes within their *Brevipalpus* vector and interact with a functionally coherent subset of mite proteins. This supports a vector-centric model in which the mite, not the plant, may be the primary adaptive environment for these viruses. Despite their distinct evolutionary histories and replication strategies, both viruses converge on molecular pathways, particularly vesicular trafficking, ER/Golgi networks, and receptor-like recognition, that likely underpin virus-vector specificity. Together, these findings support a model in which BTV glycoproteins function primarily within arthropod vectors, facilitating entry, intracellular movement, and assembly, while the plant host serves mainly as a transient transmission platform. By defining these interaction networks and validating representative components in an arthropod-like system, this study establishes a mechanistic foundation for understanding how BTV glycoproteins operate within the vector and provides a conceptual basis for future work on the determinants of transmission.

## Acknowledgments

This project was supported in part by FAPESP (Process numbers: 2017/50334-3, 2019/25078-9, 2020/15413-2, and 2023/09518-4).

