## Supplementary Figures for "Cilevirus and dichorhavirus glycoproteins target overlapping host proteins in the *Brevipalpus yothersi* vector"

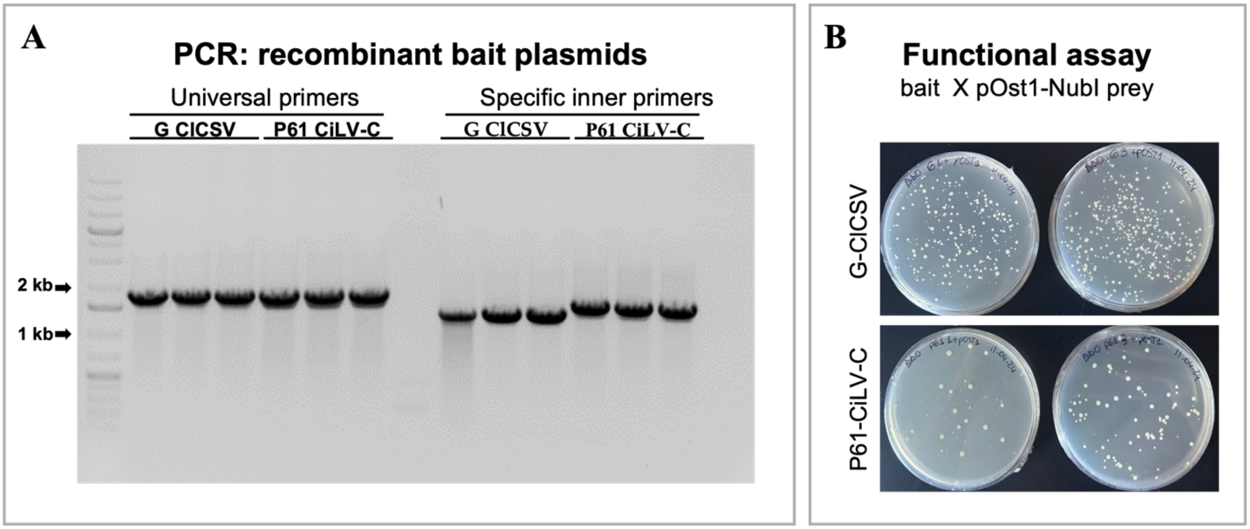


**Supplementary Figure S1. Analysis of cloning, expression, and functionality of BTV glycoproteins (P61 and G) in pBTR3-SUC bait plasmids**. **A**. 0.8% agarose gel electrophoresis showing PCR amplicons obtained from recombinant pBTR3-SUC plasmids using universal plasmid primers and insert-specific primers (Supplementary Table S1). (B) MbY2H assays showing yeast growth after 4 days on QDO selective medium following co-transformation of recombinant bait plasmids (G or P61) with the pOst1-NubI prey plasmid, with two plates per treatment.

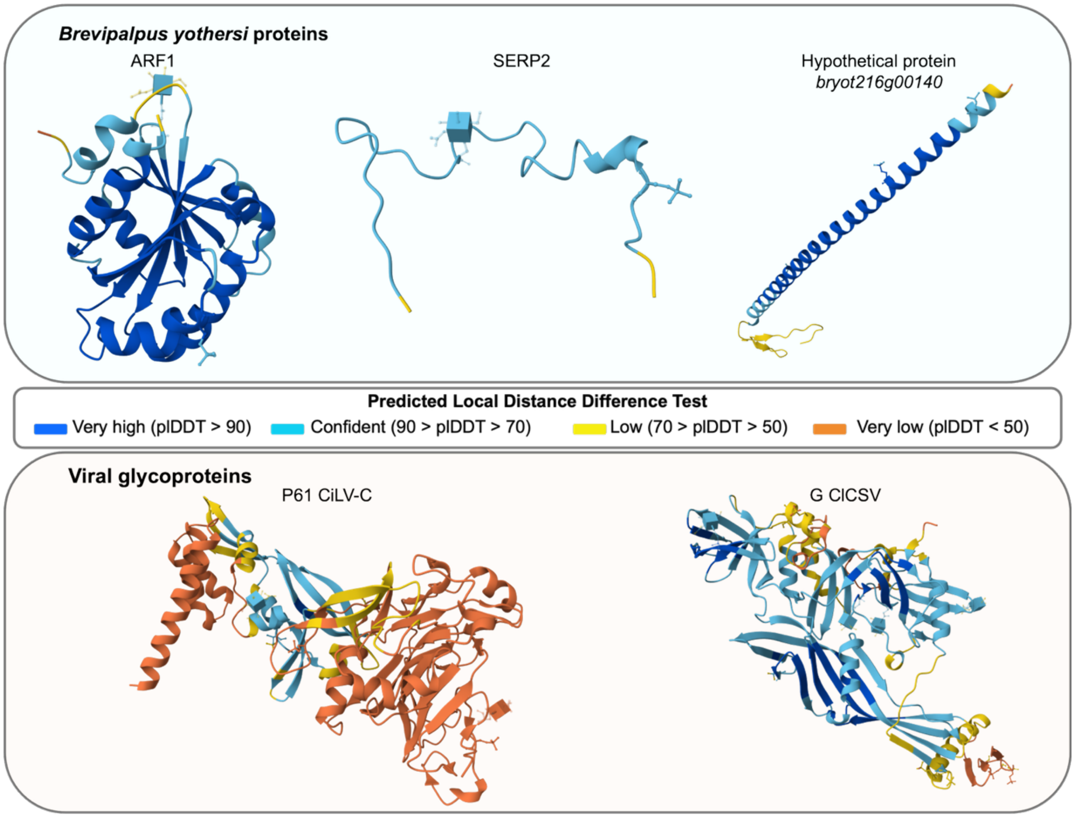


**Supplementary Figure S2. Three-dimensional structural predictions of *Brevipalpus yothersi* proteins and the viral glycoproteins P61 and G generated with AlphaFold 3.0 for protein-protein interaction analysis.** Predicted transmembrane domains and signal peptide regions were removed prior to modeling, and relevant post-translational modifications were incorporated into the final structures.

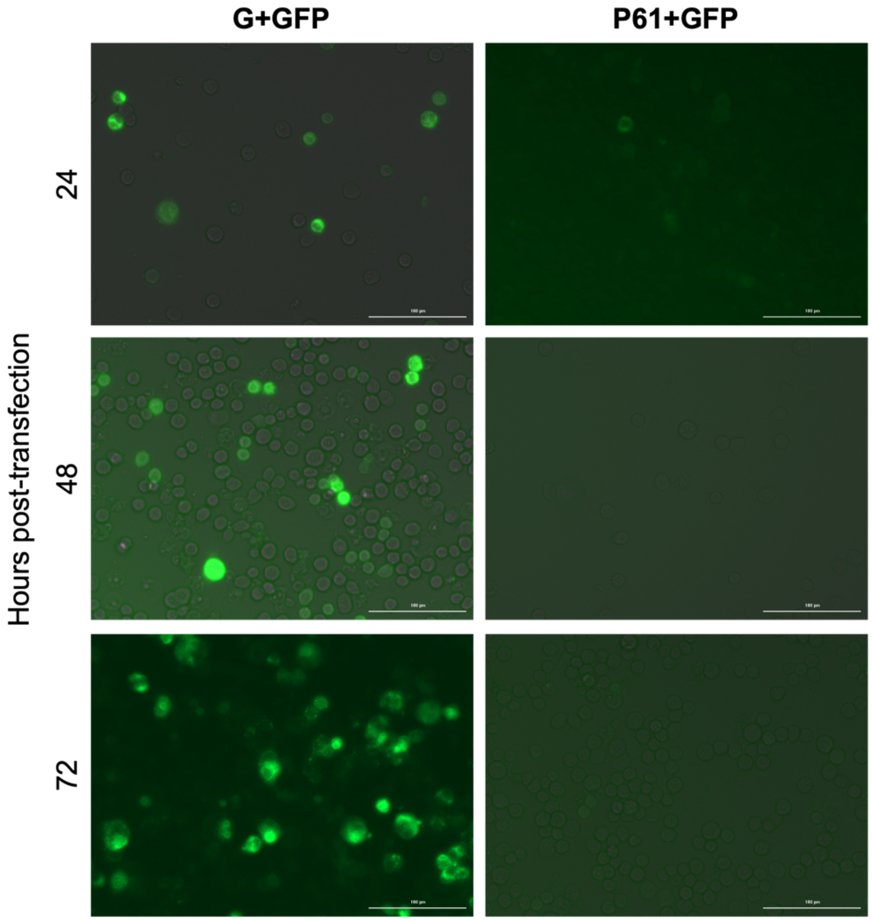


**Supplementary Figure S3. Time course of viral glycoprotein (G-ClCSV and P61-CiLV-C) transient expression in *Spodoptera frugiperda* (Sf9) cells using the pI-GFP expression vector.** Images were acquired with the Agilent BioTek imaging system at 24-, 48-, and 72-hours post-transfection.

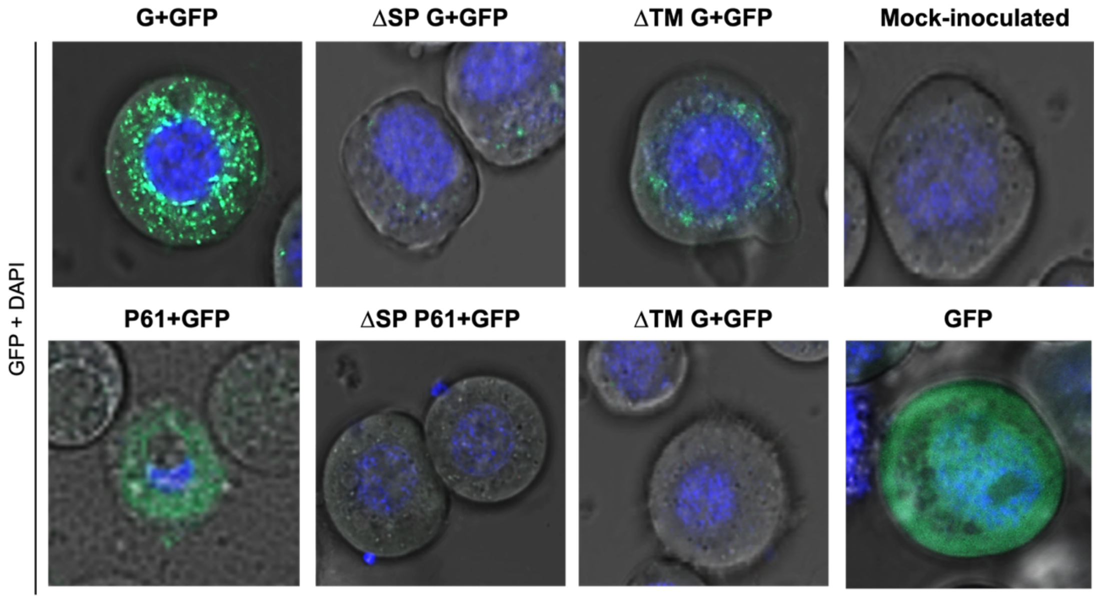


**Supplementary Figure S4. Transient expression of viral glycoproteins and their mutants in Sf9 cells using the pI-GFP expression vector.** GFP fluorescence images were acquired using a Zeiss LSM 980 confocal microscope at 72 hours post-transfection (hpt) for G-ClCSV constructs and 24 hpt for P61-CiLV-C constructs. Mutant versions were generated by deleting the predicted signal peptide (ΔSP) or the predicted transmembrane domain (ΔTM) from each glycoprotein sequence.

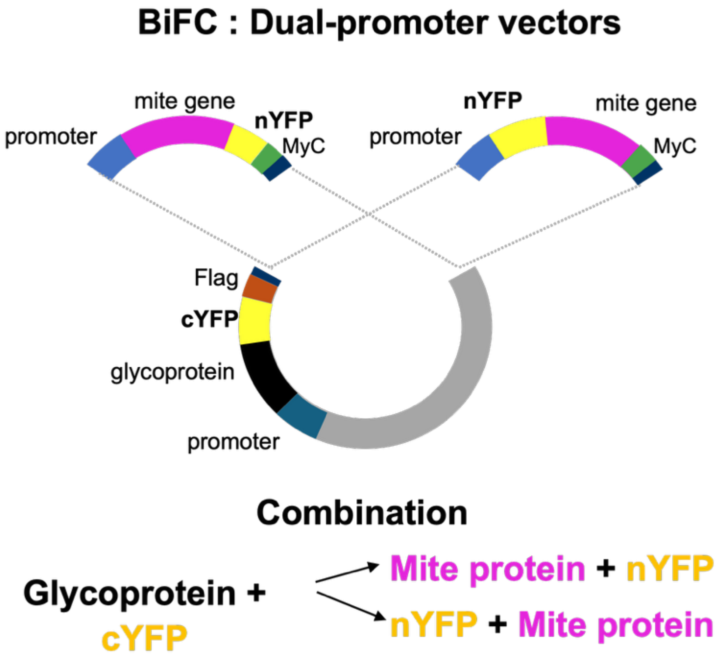


**Supplementary Figure S5. Schematic representation of the BiFC dual-promoter vector system.** The cYFP domain (yellow) was fused to the C-terminus of each viral glycoprotein (black) to generate monocistronic pIcYFP recombinant constructs. Similarly, the nYFP domain (yellow) was fused to either the N- or C-terminus of *Brevipalpus yothersi* candidate genes (pink), producing monocistronic pInYFP recombinant plasmids. For each interaction pair, the complete transcriptional unit (promoter–gene–nYFP–Myc–terminator or promoter–nYFP–gene–Myc–terminator) was PCR-amplified and inserted into the pIcYFP construct using the HiFi DNA assembly method, generating dual-promoter vectors used to assess protein–protein interactions in Sf9 cells.
