## Supplementary material for "Cilevirus and dichorhavirus glycoproteins target overlapping host proteins in the *Brevipalpus yothersi* vector": Suplementary Table S1

**Supplementary Table S1**. List of primers designed and used in this study.

| Primer name | Sequence (5′-3′) | Size (bp) | Tm (ºC) |
|---|---|---|---|
| **Bait yeast vectors** | | | |
| F_pBT3_SUC | ATTAAGGCCGCCTCGGCCATCTG | 7617 | 66 |
| R_pBT3_SUC | TAAAGGCCTGGCCGTAATGGCC | | 64 |
| F_G_bait | ACGGCCAGGCCTTTAACAGGTTGCACAAACATTCTGCC | 1575 | 71 |
| R_G_bait | GGCCGAGGCGGCCTTAATTTTCTGTTTCGCCCATGCAA | | 72 |
| F_p61_bait | ACGGCCAGGCCTTTACTTGTTTCCAATATTTACAATAGTACGGG | 1604 | 68 |
| R_p61_bait | CAGATGGCCGAGGCGGCCTTAATTAAATCAATGGTATCAGCTATGTC | | 69 |
| **PCR** | | | |
| pBT3_SUC_F_PCR | GCAAACACAAATACACACAC | - | 54 |
| pBT3_SUC_R_PCR | CTTACCGGCAAAGATCAATC | | 55 |
| pPR3-N_F_PCR | GTCGAAAATTCAAGACAAGG | - | 54 |
| pPR3-N_R_PCR | AAGCGTGACATAACTAATTAC | | 54 |
| pIW-F-PCR | TATAAATACAGCCCGCAACG | - | 55 |
| pIWRFP-R-PCR | CATGAACTCCTTGATGACGT | - | 55 |
| pIWGFP-R-PCR | GTGGTGCAGATGAACTTCAG | - | 55 |
| pIW_F_2proteins_PCR | AAAGACCCCAACGAGAAG | - | 54 |
| pIW_R_2proteins_PCR | CTTGGTCACCTTCAGCTT | | 55 |
| **BiFC empty vectors** | | | |
| R_control_pIWcYF | TCCCGGGTCGATCACTCGAGGATCCAGCGCTGGGC | - | 68 |
| F_control_pIcYWF | AGCGCTGGATCCTCGGTGATCGACCCGGGTCTAGAG | - | 67 |
| R_control_pIcYWF | CGAGGATCCAGCGCTGGGCTGCAG | - | 60 |
| R_control_pIWnYM | CGGGTCGATCACCACTCGAGGATCCAGCGCTGGGC | - | 67 |
| F_control_pInYWM | GCTGGATCCTCGATCGATCGACCCGGGTCTGAGGGCCCGCGGTTCG | - | 68 |
| **Monocistronic BiFC vectors** | | | |
| F_pIWcYF | GTGATCGACCCGGGAGGGC | - | 63 |
| R_pIW_BiFC | TCGAGGATCCAGCGCTGGG | - | 63 |
| F_pIWnYM_HiFi | GTGGTGATCGACCCGGGAG | - | 60 |
| F_pInYWM_HiFi | GATCGACCCGGGTCTGAGGGCCCGCGGTTCG | - | 68 |
| R_pInYWM_HiFi | GATCGAGGATCCAGCGCTGGG | - | 59 |
| ***B. yothersi* mite constructions - BiFC** | | | |
| F_pIW_33g00510 | GCGCTGGATCCTCGAATGAGGCAGCAAAAGATGAAAG | 471 | 57 |
| R_pIW_33g00510 | CGGGTCGATCACCACATATCCATAACGAATGCTTTGAA | | 60 |
| F_pIW_96g00450 | GCGCTGGATCCTCGAATGGCTATTGATTTAAGTGCC | 1470 | 58 |

| | | | |
|---|---|---|---|
| R_pIW_96g00450 | CGGGTCGATCACCACTACGTTGTACCAGTCC AAAAAG | | 57 |
| F_pIW_48g00250 | GCGCTGGATCCTCGAATGTCTAGCGGACGA CCTTC | 546 | 59 |
| R_pIW_48g00250 | CGGGTCGATCACCACAACTTTTTCGGTTCTTC GTC | | 61 |
| F_pIW_171g00300 | GCGCTGGATCCTCGAATGGGTAATATTGTGG CAAATC | 756 | 59 |
| R_pIW_171g00300 | CGGGTCGATCACCACTTTTTGACTYTTYAACT GATTTG | | 62 |
| F_pIW_216g00140 | GCGCTGGATCCTCGAATGATAAAAAGGCTA ACATTCA | 1089 | 60 |
| R_pIW_216g00140 | CGGGTCGATCACCACTTCAGTTTTACCCTTC GAGGTG | | 60 |
| F_pIW_217g00200 | GCGCTGGATCCTCGAATGTCAAAGGGAAAT AGTGG | 1089 | 58 |
| R_pIW_217g00200 | CGGGTCGATCACCACACAGCTACATTCTGAT TTACG | | 59 |
| F_pIW_385g00070 | GCGCTGGATCCTCGAATGGATCAGCTATTCA GTAG | 195 | 60 |
| R_pIW_385g00070 | CGGGTCGATCACCACAATGGCATCTGCAGC TTTAC | | 63 |
| F_pInY_33g00510 | GCTGGATCCTCGATCATGAGGCAGCAAAAG ATGAAAG | 471 | 60 |
| R_pInY_33g00510 | AGACCCGGGTCGATCATATCCATAACGAAT GCTTTGA | | 58 |
| F_pInY_48g00250 | GCTGGATCCTCGATCATGTCTAGCGGACGAC CTTC | 1470 | 59 |
| R_pInY_48g00250 | AGACCCGGGTCGATCAACTTTTTCGGTTCTT CGTC | | 59 |
| F_pInY_96g00450 | GCTGGATCCTCGATCATGGCTATTGATTTAA GTGCC | 546 | 61 |
| R_pInY_96g00450 | AGACCCGGGTCGATCTACGTTGTACCAGTCC AAAAAG | | 59 |
| F_pInY_171g00300 | GCTGGATCCTCGATCATGGGTAATATTGTGG CAAATC | 756 | 62 |
| R_pInY_171g00300 | AGACCCGGGTCGATCTTTTTGACTYTTYAACT GATTTG | | 60 |
| F_pInY_216g00140 | GCTGGATCCTCGATCATGATAAAAAGGCTA ACATTCA | 1089 | 60 |
| R_pInY_216g00140 | AGACCCGGGTCGATCTTCAGTTTTACCCTTC GAG | | 58 |
| F_pInY_217g00200 | GCTGGATCCTCGATCATGTCAAAGGGAAAT AGTGG | 1089 | 59 |
| R_pInY_217g00200 | AGACCCGGGTCGATCACAGCTACATTCTGAT TTAC | | 60 |
| F_pInY_385g00070 | GCTGGATCCTCGATCATGGATCAGCTATTCA GTAG | 195 | 63 |
| R_pInY_385g00070 | AGACCCGGGTCGATCAATGGCATCTGCAGC TTTAC | | 60 |
| **CiLV-C and ClCSV virus constructions - BiFC** | | | |

| | | | |
|---|---|---|---|
| F_G_pIWcYF | GCGCTGGATCCTCGAATGACCCGTTATTTGTTATCAATTC | 1575 | 63 |
| R_G_pIWcYF | TCCCGGGTCGATCACTTTCTGTTTCGCCCATGCAAC | | 60 |
| F_p61_pIWcYF | GCGCTGGATCCTCGAATGGCGCTATTTCAGCTTTTTAG | 1604 | 64 |
| R_p1_pIWcYF | TCCCGGGTCGATCACTAAATCAATGGTATCAGCTATG | | 63 |
| **Bicistronic BiFC vectors** | | | |
| F_HiFi_YFP-C | ATAACCGTATTACCGCCTTTGAG | 3638 | 57 |
| R_HiFi_YFP-C | CCACAGAATCAGGGGATAACG | | 58 |
| F_HiFi_YFP-N | CGGTAATACGGTTATCATGATGATAAACAATGTATGG | 4008 | 62 |
| R_HiFi_YFP-N | CCCCTGATTCTGTGGCACGCGCTTGAAAGGAGTGTG | | 63 |
| **Transient expression in *Sf*9 insect cells** | | | |
| **Monocistronic empty vectors** | | | |
| F_HiFi_piWG_control | ATCGACCCGGGAGGGCGCGCCATGGTGAGCAAGGGCGAGG | - | 60 |
| F_HiFi_piWR_control | ATCGACCCGGGAGGGCGCGCCATGGCCTCCTCCGAGGACGTC | - | 59 |
| R_HiFi_piW_control | CCCTCCCGGGTCGATCGATCGAGGATCCAGCGCTGGGCTGCAG | - | 61 |
| | **Monocistronic vectors** | | |
| F_pIGFP_RFP_HiFi | ATCGACCCGGGAGGGCG | 4062 | 66 |
| R_pIGFP_RFP_HiFi | CGATCGAGGATCCAGCG | | 66 |
| F_G_ClCSV_pIGFP | CGCTGGATCCTCGATCGATGACCCGTTATTTGTTATC | 1633 | 58 |
| R_G_ClCSV_pIGFP | CGCCCTCCCGGGTCGATTTTCTGTTTCGCCCATGCAAC | | 58 |
| F_p61_CiLV-C_pIGFP | CGCTGGATCCTCGATCGATGGCGCTATTTCAGCTTTTTAG | 1645 | 58 |
| R_p61_CiLV-C_pIGFP | CGCCCTCCCGGGTCGATTAAATCAATGGTATCAGCTATG | | 58 |
| F_P61_withoutSP_pIGFP | TCGATGCTTGTTTCCAATATTTACAATAGTACGGGATACTTG | 5631 | 64 |
| R_P61_ withoutSP_pIGFP | GGAAACAAGCATCGATCGAGGATCCAGCGCTGGGC | | 72 |
| F_P61_withoutTD_pIGFP | GTACTTTTTAATGCGTATATACAATTAGTGTATTTTGAGAG | 5523 | 58 |
| R_P61_ withoutTD_pIGFP | CGCATTAAAAAGTACCTTAAAGACTGTACTAAGACATTC | | 60 |
| F_G_withoutSP_pIGFP | TCGATGACAGGTTGCACAAACATTCTGCCATCAACCCTATG | ~5600 | 69 |
| R_ G _ withoutSP_pIGFP | GCAACCTGTCATCGATCGAGGATCCAGCGCTGGGCTGC | | 75 |
| F_ G _withoutTD_pIGFP | GCTTGGATCTTCAAGCTTATTATTGCAATCTTAACCAG | | 68 |
| R_ G _ withoutTD_pIGFP | CTTGAAGATCCAAGCCTTGATGTCAGGCCACTTAGATTTTAC | | 66 |
| **Recombinant and empty bicistronic vectors** | | | |
| F_Bicistronic_GFP | GGATCGATGCTCACTCAAAG | - | 64 |

| | | | |
|---|---|---|---|
| R_Bicistronic_GFP | GCCGCTACCCACGCGCTTGAAAGG | | 64 |
| F_Bicistronic_RFP | CCTTTCAAGCGCGTGGGTAGCGGCCATGATG ATAAACAATGTATG | - | 64 |
| R_Bicistronic_RFP | CTTTGAGTGAGCATCGATCC | | 64 |
| **Bicistronic P2A empty vectors** | | | |
| F_P2A_control | CTGCTGAAGCAGGCAGGCGATGTGGAGGAG AATCCCGGCCCTATGGCCTCCTCCGAGGACG TCATCAAGG | - | 70 |

**Supplementary Table S2.** Prey clones recovered from the MbY2H screens with P61-CiLV-C and G-ClCSV and summary of validation assays. Each prey plasmid recovered during the MbY2H screening was re-transformed into *E. coli* and Sanger-sequenced for confirmation. Plasmid identifiers correspond to the sequential order in which clones were collected from yeast and subsequently maintained as permanent *E. coli* stocks. Selected prey plasmids were subjected to individual MbY2H assays (each prey tested against each bait), and a subset was evaluated in duplicate under increased stringency (3-AT assays #1 and #2; 5 mM for P61 and 10 mM for G). Clones that were not tested (NT) in one or more assays are indicated in the Table.

| MbY2H screening using cDNA prey *Brevipalpus yothersi* | | | | | **Individual MbY2H assay**[1] | **3-AT assay** | |
|---|---|---|---|---|---|---|---|
| **N°** | **Interactor** | ***E. coli* plasmid #** | **Insert size-Kb** | **BLASTn and x result *B. yothersi* access ORCAE number/ gene name** | | **1** | **2** |
| 1 | G | 21 | 1.5 | bryot285g00100 Failed axon connections isoform X1 | + | 0 | 0 |
| 2 | G | 23 | 1.2 | bryot101g00380 Motile sperm domain-containing protein 1 isoform X1 | + | NT | NT |
| 3 | P61 | 237 | 1.4 | bryot101g00380 Motile sperm domain-containing protein 1 isoform X1 | + | NT | NT |
| 4 | G | 26 | 2 | bryot203g00130 Vitelline membrane outer layer protein 1 homolog | + | NT | NT |
| 5 | G | 27 | 1 | bryot34g00310 Translocon-associated protein subunit beta | + | NT | NT |
| 6 | G | 32 | 1.2 | bryot112g00030 60S ribosomal protein L5 | NT | NT | NT |
| 7 | G | 34 | 1.3 | bryot432g00010 Sodium channel protein para isoform X7 | NT | NT | NT |

| | | | | | | | |
|---|---|---|---|---|---|---|---|
| 8 | G | 55 | 1.2 | bryot150g00400<br>B-cell receptor-associated protein 31 | + | NT | NT |
| 9 | G | 41 | 1.2 | | | NT | NT |
| 10 | G | 42 | 1 | bryot107g00260<br>Tumor protein D54 isoform X3 | NT | NT | NT |
| 11 | G | 46 | 1 | bryot150g00650<br>Receptor expression-enhancing protein 6 isoform X1 | + | NT | NT |
| 12 | G | 68 | 1.1 | bryot60g00090<br>Sodium-dependent phosphate transport protein 2B | NT | NT | NT |
| 13 | G | 74 | 1.6 | bryot209g00050<br>Alpha-(1,3)-fucosyltransferase C-like | + | NT | NT |
| 14 | G | 79 | 0.9 | bryot96g00450<br>Syntaxin-18 | + | 3 | 3 |
| 15 | G | 92 | 1.6 | bryot194g00080<br>Acyl-CoA desaturase | + | 3 | 3 |
| 16 | G | 95 | 1 | bryot11g00420<br>Thioredoxin-like protein 1 | NT | NT | NT |
| 17 | G | 104 | 1 | bryot05g01040<br>Geranylgeranyl transferase type-2 subunit beta | NT | NT | NT |
| 18 | G | 110 | 1.1 | bryot312g00040<br>ATP synthase subunit d, mitochondrial | NT | NT | NT |
| 19 | G | 493 | 1 | | NT | NT | NT |
| 20 | G | 119 | 0.8 | bryot370g00030<br>putative ATP-dependent RNA helicase TDRD9 | NT | NT | NT |
| 21 | G | 125 | 1.2 | bryot48g00250<br>40S ribosomal protein SA | + | 3 | 3 |
| 22 | G | 346 | 1.2 | | | | |
| 23 | G | 646 | 1.4 | | | | |
| 24 | G | 847 | 1.2 | | | | |
| 25 | P61 | 31 | 1.3 | bryot48g00250<br>40S ribosomal protein SA | + | 0 | 0 |
| 26 | G | 128 | 0.9 | bryot207g00090<br>Nuclear receptor coactivator 6-like | NT | NT | NT |

| | | | | | | | |
|---|---|---|---|---|---|---|---|
| 27 | G | 133 | 1 | bryot84g00020 Translocon-associated protein subunit gamma | NT | NT | NT |
| 28 | G | 122 | 1 | | | | |
| 29 | G | 43 | 1 | | | | |
| 30 | G | 137 | 1.1 | bryot265g00150 Tyrosine-protein phosphatase non-receptor type 23 | NT | NT | NT |
| 31 | G | 147 | 1.1 | bryot171g00300 ADP-ribosylation factor 1 | + | 3 | 3 |
| 32 | G | 385 | 1.1 | | | | |
| 33 | P61 | 25 | 1.6 | bryot171g00300 ADP-ribosylation factor 1 | + | 1 | 1 |
| 34 | P61 | 122 | 1.2 | | | | |
| 35 | P61 | 217 | 1.4 | | | | |
| 36 | P61 | 269 | 1.4 | | | | |
| 37 | P61 | 299 | 1.2 | | | | |
| 38 | P61 | 355 | 1.5 | | | | |
| 39 | P61 | 390 | 1.2 | | | | |
| 40 | P61 | 404 | 1.3 | | | | |
| 41 | G | 149 | 1 | bryot216g00140 Hypothetical protein | + | 3 | 3 |
| 42 | G | 193 | 0.9 | | | | |
| 43 | P61 | 370 | 0.9 | bryot216g00140 Hypothetical protein | + | 2 | 2 |
| 44 | G | 151 | 1.2 | bryot33g00510 Stress-associated endoplasmic reticulum protein 2 | + | 3 | 3 |
| 45 | G | 457 | 0.9 | | | | |
| 46 | P61 | 113 | 0.9 | bryot33g00510 Stress-associated endoplasmic reticulum protein 2 | + | 1 | 1 |
| 47 | G | 152 | 1.2 | bryot412g00080 Ras-related protein Rab-18 | + | NT | NT |

| | | | | | | | |
|---|---|---|---|---|---|---|---|
| 48 | G | 157 | 1.3 | bryot26g00390 Transmembrane emp24 domain-containing protein 5-like | + | NT | NT |
| 49 | G | 158 | 0.8 | bryot471g00040 Hypothetical protein | + | 2 | 2 |
| 50 | G | 161 | 1 | bryot139g00170 uncharacterized protein LOC107366498 | + | 3 | 3 |
| 51 | G | 166 | 1 | bryot98g00340 RING-box protein 1a | NT | NT | NT |
| 52 | G | 174 | 1.2 | bryot580g00010 Hypothetical protein | + | 0 | 0 |
| 53 | G | 208 | 1 | | | | |
| 54 | G | 296 | 1.2 | | | | |
| 55 | G | 370 | 1 | | | | |
| 56 | G | 384 | 1 | | | | |
| 57 | G | 442 | 1 | | | | |
| 58 | G | 494 | 1.1 | | | | |
| 59 | G | 608 | 1.1 | | | | |
| 60 | G | 686 | 1.2 | | | | |
| 61 | P61 | 120 | 1.1 | bryot580g00010 Hypothetical protein | + | 0 | 0 |
| 62 | G | 177 | 1.2 | bryot33g00070 60S ribosomal protein L23a | NT | NT | NT |
| 63 | G | 178 | 1 | bryot02g01480 Gamma-aminobutyric acid receptor-associated protein | + | 3 | 3 |
| 64 | G | 185 | 1 | | | | |
| 65 | G | 272 | 1.1 | | | | |
| 66 | G | 394 | 1 | | | | |
| 67 | G | 397 | 1 | | | | |

| | | | | | | | |
|---|---|---|---|---|---|---|---|
| 68 | G | 503 | 1 | | | | |
| 69 | P61 | 158 | 1 | bryot02g01480 Gamma-aminobutyric acid receptor-associated protein | + | NT | NT |
| 70 | G | 180 | 1.2 | bryot05g00800 Piwi-like protein 1 | NT | NT | NT |
| 71 | G | 187 | 1.1 | bryot63g00220 60S ribosomal protein L10 | NT | NT | NT |
| 72 | G | 885 | 1.1 | | | | |
| 73 | G | 460 | 1 | | | | |
| 74 | P61 | 13 | 1.1 | bryot63g00220 60S ribosomal protein L10 | NT | NT | NT |
| 75 | P61 | 152 | 1.1 | | | | |
| 76 | G | 195 | 1.2 | bryot05g00920 T-complex protein 1 subunit epsilon | NT | NT | NT |
| 77 | G | 196 | 2 | bryot23g00510 Uncharacterized protein CG7065 | + | NT | NT |
| 78 | G | 525 | 1.2 | | | | |
| 79 | G | 263 | 1.6 | | | | |
| 80 | P61 | 236 | 1.6 | bryot23g00510 Uncharacterized protein CG7065 | + | 3 | 3 |
| 81 | G | 216 | 1 | bryot03g01070 Translocation protein SEC62 | + | NT | NT |
| 82 | G | 218 | 1 | | | NT | NT |
| 83 | G | 219 | 1.1 | bryot407g00060 60S ribosomal protein L7a | NT | NT | NT |
| 84 | G | 358 | 1 | | | | |
| 85 | P61 | 55 | 1.2 | bryot407g00060 60S ribosomal protein L7a | NT | NT | NT |
| 86 | G | 220 | 1.1 | bryot27g00630 protein catecholamines up | + | NT | NT |
| 87 | G | 882 | 1.2 | | | NT | NT |

| | | | | | | | |
|---|---|---|---|---|---|---|---|
| 88 | G | 224 | 1.1 | bryot401g00010 PAX-interacting protein 1 | NT | NT | NT |
| 89 | G | 226 | 1.2 | | | NT | NT |
| 90 | G | 225 | 1.3 | bryot200g00330 Myosin regulatory light polypeptide 9 | NT | NT | NT |
| 91 | G | 231 | 1.2 | bryot219g00190 Glyceraldehyde-3-phosphate dehydrogenase | + | NT | NT |
| 92 | G | 232 | 1 | bryot505g00010 Hypothetical protein | NT | NT | NT |
| 93 | G | 238 | 1.1 | bryot108g00160 Hypothetical protein | + | NT | NT |
| 94 | G | 241 | 0.9 | bryot185g00040 ATP synthase lipid-binding protein, mitochondrial | + | NT | NT |
| 95 | G | 245 | 1.5 | bryot285g00040 Peroxidase | + | NT | NT |
| 96 | G | 248 | 0.9 | bryot10g00330 Hypothetical protein | + | 2 | 2 |
| 97 | G | 415 | 1 | | | | |
| 98 | P61 | 430 | 0.9 | bryot10g00330 Hypothetical protein | + | 0 | 0 |
| 99 | G | 252 | 2 | bryot119g00040 Elongation of very long chain fatty acids protein AAEL008004-like | + | 0 | 0 |
| 100 | G | 254 | 1 | bryot376g00150 R3H domain-containing protein 1-like isoform X4 | NT | NT | NT |
| 101 | P61 | 44 | 0.9 | bryot376g00150 R3H domain-containing protein 1-like isoform X4 | NT | NT | NT |
| 102 | P61 | 59 | 1 | | | | |
| 103 | P61 | 212 | 0.9 | | | | |
| 104 | P61 | 278 | 1.1 | | | | |
| 105 | P61 | 308 | 0.9 | | | | |
| 106 | P61 | 322 | 2.1 | | | | |
| 107 | P61 | 420 | 0.9 | | | | |

| | | | | | | | |
|---|---|---|---|---|---|---|---|
| 108 | G | 260 | 0.6 | bryot235g00080 Chondroitin proteoglycan-2-like | NT | NT | NT |
| 109 | G | 266 | 1.4 | bryot13g00270 Signal transducing adapter molecule 1 | + | NT | NT |
| 110 | G | 268 | 1.5 | bryot143g00270 Disintegrin and metalloproteinase domain-containing protein 15 isoform X1 | + | NT | NT |
| 111 | G | 275 | 1.2 | bryot386g00040 Surfeit locus protein 4 homolog | + | NT | NT |
| 112 | G | 284 | 1.1 | bryot27g00270 Magnesium-dependent phosphatase 1-like | + | NT | NT |
| 113 | G | 288 | 1.2 | bryot41g00470 Protein spinster homolog 1 isoform X2 | + | NT | NT |
| 114 | G | 289 | 2 | bryot284g00070 Nucleosome assembly protein 1-like 4, partial | NT | NT | NT |
| 115 | G | 290 | 1 | bryot678g00020 Leucine-rich repeat neuronal protein 1-like | + | 1 | 1 |
| 116 | G | 308 | 1.2 | | | | |
| 117 | G | 522 | 1 | | | | |
| 118 | G | 594 | 0.8 | | | | |
| 119 | G | 771 | 0.9 | | | | |
| 120 | G | 773 | 1 | | | | |
| 121 | G | 795 | 1 | | | | |
| 122 | G | 298 | 1.6 | bryot22g00740 Ribosome biogenesis protein BRX1 homolog | NT | NT | NT |
| 123 | G | 302 | 0.8 | bryot436g00070 Hypothetical protein | NT | NT | NT |
| 124 | G | 305 | 0.9 | bryot223g00030 63 kDa sperm flagellar membrane protein | NT | NT | NT |
| 125 | G | 310 | 1.3 | bryot435g00040 Protein regulator of cytokinesis 1 isoform X1 | NT | NT | NT |

| | | | | | | | |
|---|---|---|---|---|---|---|---|
| 126 | G | 320 | 1.8 | bryot325g00040 Ubiquinone biosynthesis O-methyltransferase, mitochondrial | + | NT | NT |
| 127 | G | 322 | 1.2 | bryot385g00080 Zinc finger protein 512B-like | NT | NT | NT |
| 128 | G | 323 | 1 | bryot317g00050 Putative ATP-dependent RNA helicase me31b | NT | NT | NT |
| 129 | G | 562 | 1 | | | NT | NT |
| 130 | G | 328 | 1 | bryot02g00990 Mitogen-activated protein kinase kinase kinase kinase 5 isoform X1 | NT | NT | NT |
| 131 | G | 331 | 1.1 | bryot72g00150 Presqualene diphosphate phosphatase | + | NT | NT |
| 132 | G | 334 | 1 | bryot634g00040 40S ribosomal protein S9 | NT | NT | NT |
| 133 | G | 340 | 1.2 | bryot13g00880 60S ribosomal protein L8 | NT | NT | NT |
| 134 | G | 349 | 2 | bryot296g00020 eukaryotic translation initiation factor 5 | NT | NT | NT |
| 135 | G | 352 | 1.00 | bryot613g00010 Dihydropyrimidinase-like | NT | NT | NT |
| 136 | G | 354 | 2 | bryot18g00240 Dolichyl-diphosphooligosaccharide--protein glycosyltransferase subunit 2 isoform X1 | + | NT | NT |
| 137 | G | 355 | 1 | bryot11g00600 Serum response factor | NT | NT | NT |
| 138 | G | 363 | 1 | bryot72g00270 Hypothetical protein X777_15795 | + | 3 | 3 |
| 139 | G | 545 | 1.1 | | | | |
| 140 | G | 888 | 1.2 | | | | |
| 141 | P61 | 1 | 1.03 | bryot72g00270 Hypothetical protein X777_15795 | + | 0 | 0 |
| 142 | P61 | 2 | 1.05 | | | | |
| 143 | P61 | 3 | 1.1 | | | | |

| | | | | | | | |
|---|---|---|---|---|---|---|---|
| 144 | P61 | 30 | 1.1 | | | | |
| 145 | P61 | 76 | 1.4 | | | | |
| 146 | P61 | 109 | 1.4 | | | | |
| 147 | P61 | 276 | 1.3 | | | | |
| 148 | G | 373 | 1 | bryot64g00230 Neuroligin-4, X-linked-like, partial | + | 2 | 3 |
| 149 | G | 376 | 1.2 | bryot05g00810 Ribosome maturation protein SBDS | NT | NT | NT |
| 150 | G | 379 | 1.2 | bryot281g00100 ER lumen protein-retaining receptor 2 | + | NT | NT |
| 151 | G | 598 | 1.5 | | | NT | NT |
| 152 | G | 736 | 1.5 | | | NT | NT |
| 153 | G | 382 | 1 | bryot86g00350 N-alpha-acetyltransferase 30 | NT | NT | NT |
| 154 | G | 534 | 1 | | | NT | NT |
| 155 | G | 386 | 1.2 | bryot35g00670 Chromosome segregation protein SMC | NT | NT | NT |
| 156 | G | 388 | 2 | bryot201g00160 Zinc transporter 9 | + | NT | NT |
| 157 | G | 409 | 1 | bryot37g00020 Hypothetical protein | + | NT | NT |
| 158 | G | 412 | 1.4 | bryot584g00070 COP9 signalosome complex subunit 2 | NT | NT | NT |
| 159 | G | 421 | 1.4 | bryot137g00190 Hypothetical protein | + | NT | NT |
| 160 | G | 424 | 0.9 | bryot281g00020 Sodium/myo-inositol cotransporter 2 | NT | NT | NT |
| 161 | G | 433 | 1.2 | bryot295g00050 DNA topoisomerase 2-alpha | NT | NT | NT |
| 162 | G | 445 | 1 | bryot32g00120 Mitochondrial inner membrane protein OXA1L | + | NT | NT |

| | | | | | | | |
|---|---|---|---|---|---|---|---|
| 163 | G | 463 | 1.2 | bryot391g00080<br>Motile sperm domain-containing protein 2-like | + | NT | NT |
| 164 | G | 470 | 1 | bryot267g00170<br>Hypothetical protein | + | NT | NT |
| 165 | G | 478 | 1 | bryot150g00170<br>elongation of very long chain fatty acids protein AAEL008004-like | + | NT | NT |
| 166 | P61 | 70 | 1 | bryot150g00170<br>elongation of very long chain fatty acids protein AAEL008004-like | + | 2 | 3 |
| 167 | G | 479 | 1.2 | bryot92g00220<br>Carboxypeptidase E | + | NT | NT |
| 168 | G | 481 | 1 | bryot151g00190<br>AP-2 complex subunit sigma | + | NT | NT |
| 169 | G | 483 | 1 | bryot111g00360<br>Ribosomal protein S5, partial | NT | NT | NT |
| 170 | G | 487 | 1.5 | bryot01g00440<br>Choline kinase alpha-like isoform X2 | + | NT | NT |
| 171 | G | 491 | 1.1 | bryot49g00320<br>Hypothetical protein | + | 0 | 0 |
| 172 | G | 805 | 1.1 | | | | |
| 173 | G | 517 | 0.6 | bryot35g00310<br>Signal peptidase complex subunit 1 | + | NT | NT |
| 174 | G | 528 | 1 | bryot75g00320<br>Abhydrolase domain-containing protein 16A | NT | NT | NT |
| 175 | G | 531 | 1 | bryot201g00040<br>Uncharacterized protein LOC107359576 | + | NT | NT |
| 176 | G | 540 | 1 | bryot219g00060<br>Epididymal secretory protein E1-like | + | NT | NT |
| 177 | G | 543 | 1.2 | bryot159g00360<br>Polyubiquitin-B | + | NT | NT |
| 178 | G | 546 | 1.2 | bryot571g00080<br>Persulfide dioxygenase ETHE1, mitochondrial | NT | NT | NT |
| 179 | G | 549 | 1 | bryot57g00400<br>Uncharacterized protein LOC107360510 | NT | NT | NT |
| 180 | G | 559 | 1 | bryot22g00300<br>Clathrin light chain B-like isoform X2 | + | NT | NT |

| | | | | | | | |
|---|---|---|---|---|---|---|---|
| 181 | G | 563 | 1.1 | bryot14g00390<br>Hypothetical protein | + | 0 | 0 |
| 182 | G | 584 | 0.5-1 | bryot23g00330<br>CD63 antigen | + | NT | NT |
| 183 | G | 592 | 0.9 | bryot02g01390<br>Nose resistant to fluoxetine protein 6-like | NT | NT | NT |
| 184 | G | 607 | 1 | bryot444g00110<br>60S ribosomal protein L13a | NT | NT | NT |
| 185 | G | 627 | 1.4 | bryot274g00130<br>Eukaryotic translation initiation factor 3 subunit F | NT | NT | NT |
| 186 | G | 639 | 1.4 | bryot452g00010<br>Ribosome biogenesis protein NSA2 homolog | NT | NT | NT |
| 187 | G | 643 | 1.4 | bryot67g00380<br>Huntingtin-interacting protein K | NT | NT | NT |
| 188 | G | 655 | 1.5 | bryot36g00610<br>Dual specificity protein phosphatase 3 isoform X2 | NT | NT | NT |
| 189 | G | 658 | 1.2 | bryot29g00220<br>5-methyltetrahydropteroyltriglutamate--homocysteine S-methyltransferase | NT | NT | NT |
| 190 | G | 670 | 1.2 | bryot75g00020<br>Protein phosphatase 1 regulatory subunit 7 | NT | NT | NT |
| 191 | G | 673 | 1.2 | bryot267g00180<br>Hypothetical protein | + | 2 | 2 |
| 192 | G | 676 | 0.9 | bryot151g00220<br>Transmembrane protein 245 isoform X1 | NT | NT | NT |
| 193 | G | 680 | 1.2 | bryot27g00200<br>Proteasome activator complex subunit 3 | NT | NT | NT |
| 194 | G | 682 | 1.2 | bryot442g00050<br>Uncharacterized protein LOC107369187 | + | 3 | 3 |
| 195 | G | 688 | 1.3 | bryot257g00080<br>Eukaryotic translation initiation factor 6 | NT | NT | NT |
| 196 | G | 693 | 1 | bryot197g00280<br>Minor histocompatibility antigen H13 | + | NT | NT |
| 197 | G | 697 | 1 | | | | |
| 198 | G | 695 | 1 | bryot71g00470<br>Hypothetical protein | + | 0 | 0 |

| | | | | | | | |
|---|---|---|---|---|---|---|---|
| 199 | G | 696 | 1.1 | | | | |
| 200 | G | 709 | 1.5 | bryot217g00200<br>Ras-related protein Rab-14 isoform X1 | + | 3 | 3 |
| 201 | P61 | 457 | 1.7 | bryot217g00200<br>Ras-related protein Rab-14 isoform X1 | + | 3 | 3 |
| 202 | G | 724 | 1 | bryot72g00400<br>SEC14-like protein 1 | + | 3 | 3 |
| 203 | P61 | 11 | 1.2 | bryot72g00400<br>SEC14-like protein 1 | + | 0 | 0 |
| 204 | P61 | 12 | 1.2 | | | | |
| 205 | G | 730 | 0.8 | bryot235g00020<br>Uncharacterized protein LOC107366944 | NT | NT | NT |
| 206 | G | 753 | 1.5 | bryot335g00090<br>ATP synthase subunit gamma, mitochondrial isoform X1 | + | NT | NT |
| 207 | G | 775 | 1 | bryot126g00240<br>40S ribosomal protein S20 | NT | NT | NT |
| 208 | G | 785 | 1 | bryot43g00370<br>Hypothetical protein | + | 3 | 3 |
| 209 | G | 800 | 0.9 | bryot144g00190<br>Sphingosine-1-phosphate phosphatase 1 | + | 3 | 3 |
| 210 | G | 859 | 0.9 | | | | |
| 211 | G | 865 | 0.9 | | | | |
| 212 | G | 867 | 0.9 | | | | |
| 213 | G | 849 | 0.9 | | | | |
| 214 | G | 852 | 0.9 | | | | |
| 215 | G | 810 | 1.2 | bryot59g00170<br>Histone H2AX | NT | NT | NT |
| 216 | G | 844 | 0.9 | | | NT | NT |
| 217 | G | 817 | 1 | bryot25g00440<br>50S ribosomal protein L20 | NT | NT | NT |
| 218 | G | 818 | 1.3 | bryot238g00010<br>Syntaxin-1A isoform X2 | NT | NT | NT |

| | | | | | | | |
|---|---|---|---|---|---|---|---|
| 219 | G | 824 | 1.1 | bryot202g00100 Spectrin beta chain isoform X1 | + | NT | NT |
| 220 | G | 828 | 1 | bryot07g00430 Anamorsin | NT | NT | NT |
| 221 | G | 831 | 1 | bryot45g00160 bax inhibitor 1 | + | 3 | 3 |
| 222 | G | 835 | 1.3 | bryot45g00160 bax inhibitor 1 | + | 3 | 3 |
| 223 | G | 851 | 1.5 | bryot22g00310 Proline dehydrogenase 1, mitochondrial-like | NT | NT | NT |
| 224 | G | 856 | 1.1 | bryot01g01180 Reticulon-4 isoform X2 | + | NT | NT |
| 225 | G | 877 | 1.1 | bryot150g00200 Peroxiredoxin-1 | NT | NT | NT |
| 226 | G | 893 | 0.9 | bryot161g00070 Putative RNA-binding protein Luc7-like 2 isoform X1 | NT | NT | NT |
| 227 | P61 | 4 | 1.2 | bryot77g00200 Adult-specific rigid cuticular protein 15.7-like | + | NT | NT |
| 228 | P61 | 7 | 1.1 | bryot147g00090 tRNA-splicing endonuclease subunit Sen2 | NT | NT | NT |
| 229 | P61 | 137 | 1 | bryot147g00090 tRNA-splicing endonuclease subunit Sen2 | NT | NT | NT |
| 230 | P61 | 20 | 1.1 | bryot242g00030 V-type proton ATPase 16 kDa proteolipid subunit | + | NT | NT |
| 231 | P61 | 29 | 1.1 | bryot385g00070 Innexin inx1 | + | 3 | 2 |
| 232 | P61 | 42 | 1 | bryot385g00070 Innexin inx1 | + | 3 | 2 |
| 233 | P61 | 134 | 0.9 | bryot385g00070 Innexin inx1 | + | 3 | 2 |
| 234 | P61 | 250 | 1.4 | bryot385g00070 Innexin inx1 | + | 3 | 2 |
| 235 | P61 | 340 | 1.1 | bryot385g00070 Innexin inx1 | + | 3 | 2 |
| 236 | P61 | 39 | 1.4 | bryot173g00150 Prostate stem cell antigen | + | NT | NT |
| 237 | G | 712 | 1 | bryot745g00010 cytochrome oxidase subunit I | + | 3 | 3 |
| 238 | G | 739 | 1 | bryot745g00010 cytochrome oxidase subunit I | + | 3 | 3 |

| | | | | | | | |
|---|---|---|---|---|---|---|---|
| 239 | P61 | 72 | 1 | bryot745g00010 Cytochrome oxidase subunit I | + | 0 | 0 |
| 240 | P61 | 74 | 1 | | | | |
| 241 | P61 | 77 | 1.1 | bryot150g00410 Ethanolamine phosphotransferase 1 | + | NT | NT |
| 242 | P61 | 82 | 1 | bryot354g00190 Zinc transporter ZIP3-like | NT | NT | NT |
| 243 | P61 | 87 | 1.4 | bryot434g00010 Putative sodium-coupled neutral amino acid transporter 7 | + | NT | NT |
| 244 | P61 | 90 | 0.9—1.6 | bryot82g00160 V-type proton ATPase 16 kDa proteolipid subunit | + | 3 | 3 |
| 245 | P61 | 93 | 1.2 | bryot506g00020 Serine incorporator 1 | + | NT | NT |
| 246 | P61 | 97 | 1.1 | bryot182g00250 Zinc finger protein 40 | NT | NT | NT |
| 247 | P61 | 100 | 1.5 | bryot105g00040 Deoxyribonuclease-2-alpha | NT | NT | NT |
| 248 | P61 | 101 | 2 | bryot91g00080 Polyadenylate-binding protein-interacting protein 1 isoform X2 | NT | NT | NT |
| 249 | P61 | 103 | 1.2 | bryot630g00010 Protein odd-skipped-related 2 isoform X1 | NT | NT | NT |
| 250 | P61 | 116 | 1.1 | bryot19g00020 Peroxisomal biogenesis factor 16 | + | NT | NT |
| 251 | P61 | 119 | 1 | bryot257g00140 Long-chain fatty acid transport protein 4 | NT | NT | NT |
| 252 | P61 | 121 | 1.4 | bryot216g00190 Rab proteins geranylgeranyltransferase component A | NT | NT | NT |
| 253 | P61 | 131 | 1.1 | bryot11g00120 Hypothetical protein | + | 1 | 1 |
| 254 | P61 | 229 | 1.5 | bryot36g00730 E3 ubiquitin-protein ligase MARCH6 | + | NT | NT |
| 255 | P61 | 149 | 1.6 | | | | |
| 256 | P61 | 164 | 0.8—1.6 | bryot462g00050 E3 ubiquitin-protein ligase NRDP1, partial | NT | NT | NT |

| | | | | | | | |
|---|---|---|---|---|---|---|---|
| 257 | P61 | 173 | 1 | bryot169g00230<br>Reticulon-1 isoform X3 | + | NT | NT |
| 258 | P61 | 176 | 1.1 | bryot103g00240<br>ER membrane protein complex subunit 4 | + | 2 | 1 |
| 259 | P61 | 179 | 1.2 | bryot59g00390<br>Cyclin-dependent kinase 6 | NT | NT | NT |
| 260 | P61 | 296 | 1.7 | | | | |
| 261 | P61 | 184 | 1 | bryot68g00300<br>Hypothetical protein | + | NT | NT |
| 262 | P61 | 203 | 2 | bryot27g00500<br>Sterol O-acyltransferase 1 | + | NT | NT |
| 263 | P61 | 240 | 1 | bryot43g00190<br>Niemann-Pick C1-like protein 1 | NT | NT | NT |
| 264 | P61 | 346 | 1 | | | | |
| 265 | P61 | 243 | 1.6 | bryot127g00010<br>5'-AMP-activated protein kinase subunit beta-1 | NT | NT | NT |
| 266 | P61 | 260 | 1.1 | bryot442g00030<br>Gamma-interferon-inducible lysosomal thiol reductase-like | + | NT | NT |
| 267 | P61 | 263 | 1.2 | bryot153g00020<br>Saccharopine dehydrogenase-like oxidoreductase | + | NT | NT |
| 268 | P61 | 266 | 1.5 | bryot261g00140<br>Hypothetical protein X975_09073 | + | 0 | 0 |
| 269 | P61 | 273 | 0.8 | bryot87g00040<br>DnaJ homolog subfamily A member 3, mitochondrial | + | NT | NT |
| 270 | P61 | 275 | 1.1 | bryot114g00040<br>Cuticular protein | + | NT | NT |
| 271 | P61 | 392 | 1 | | | | |
| 272 | P61 | 287 | 1.4 | bryot18g00860<br>Long wavelength rhodopsin | + | NT | NT |
| 273 | P61 | 293 | 1.1 | bryot354g00030<br>Hypothetical protein | + | 1 | 1 |
| 274 | P61 | 304 | 1 | bryot267g00090<br>ABC transporter G family member 20 isoform X1 | + | 3 | 2 |
| 275 | P61 | 332 | 1 | | + | NT | NT |

| | | | | | | | |
|---|---|---|---|---|---|---|---|
| 276 | P61 | 360 | 1 | bryot06g00910<br>Translocating chain-associated membrane protein 1 | | | |
| 277 | P61 | 337 | 1.2 | bryot168g00180<br>transmembrane protease serine 9-like | + | NT | NT |
| 278 | P61 | 348 | 1 | bryot268g00020<br>Coatomer subunit zeta-1 | + | NT | NT |
| 279 | P61 | 350 | 2 | bryot119g00020<br>Golgi reassembly-stacking protein 2 | + | NT | NT |
| 280 | P61 | 375 | 1.7 | bryot04g00440<br>Ras-related protein Rab-8A | NT | NT | NT |
| 281 | P61 | 379 | 0.9 | bryot144g00390<br>Hypothetical protein | + | 3 | 3 |
| 282 | P61 | 393 | 1.4 | bryot224g00240<br>pre-mRNA-splicing factor SYF1 | NT | NT | NT |
| 283 | P61 | 397 | 0.9 | bryot106g00120<br>actin cytoskeleton-regulatory complex protein PAN1-like isoform X1 | NT | NT | NT |
| 284 | P61 | 413 | 1.6 | bryot148g00080<br>Ras-related protein Rab-27A | + | NT | NT |
| 285 | P61 | 424 | 1.4 | bryot126g00030<br>Nucleolar protein 10 | NT | NT | NT |
| 286 | P61 | 439 | 0.9 | bryot107g00180<br>Sodium/potassium-transporting ATPase subunit alpha isoform X1 | NT | NT | NT |
| 287 | P61 | 460 | 1.6 | bryot387g00060<br>Surfeit locus protein 4 | + | NT | NT |
| **Number** | **Interactor** | **E. coli plasmid #** | **Insert size-Kb** | **BLASTn and x result accession number/*B. obovatus* gene name** | **Individual MbY2H assay** | **3-AT assay** | |
| | | | | | | **1** | **2** |
| 1 | G | 49 | 1 | LOC141858461<br>Death-associated protein 1-like | NT | NT | NT |
| 2 | G | 86 | 1.5 | LOC141855195<br>Calcium channel flower-like | NT | NT | NT |
| 3 | G | 101 | 1.1 | LOC141849406<br>G1/S-specific cyclin-D2-like | NT | NT | NT |
| 4 | G | 121 | 1.5 | LOC141849449<br>Uncharacterized protein | NT | NT | NT |
| 5 | G | 172 | 1 | LOC141854812<br>Uncharacterized protein | NT | NT | NT |

| | | | | | | | |
|---|---|---|---|---|---|---|---|
| 6 | G | 183 | 0.9 | LOC141855901 Uuncharacterized protein | NT | NT | NT |
| 7 | G | 361 | 1 | | NT | NT | NT |
| 8 | G | 631 | 1 | | NT | NT | NT |
| 9 | G | 634 | 1 | | NT | NT | NT |
| 10 | G | 685 | 1 | | NT | NT | NT |
| 11 | G | 735 | 1 | | NT | NT | NT |
| 12 | G | 825 | 1 | | NT | NT | NT |
| 13 | G | 868 | 1 | | NT | NT | NT |
| 14 | P61 | 27 | 1.2 | LOC141855901 Uncharacterized protein | NT | NT | NT |
| 15 | P61 | 221 | 1.2 | | NT | NT | NT |
| 16 | G | 259 | 1.4 | LOC141858526 Ectonucleoside triphosphate diphosphohydrolase 3-like | NT | NT | NT |
| 17 | G | 342 | 1 | Brevipalpus obovatus transcription factor E2F3-like (LOC141853535) | NT | NT | NT |
| 18 | G | 350 | 1 | LOC141855407 uncharacterized protein | NT | NT | NT |
| 19 | G | 779 | 1 | | NT | NT | NT |
| 20 | G | 511 | 1 | LOC141857809 Transcription factor kayak-like | NT | NT | NT |
| 21 | G | 761 | 1 | | NT | NT | NT |
| 22 | G | 564 | 1 | LOC141857989 Stomatin-4-like | NT | NT | NT |
| 23 | G | 640 | 1 | LOC141848925 Uncharacterized protein | NT | NT | NT |
| 24 | G | 691 | 1 | LOC141853891 Protein dj-1beta-like | NT | NT | NT |
| 25 | G | 718 | 1 | LOC141849033 Innexin inx2-like | NT | NT | NT |
| 26 | G | 727 | 1 | Regulator of G-protein signaling 7 (RSG7) | NT | NT | NT |

| | | | | | | | |
|---|---|---|---|---|---|---|---|
| 27 | G | 807 | 1 | LOC141849144 Uncharacterized protein | NT | NT | NT |
| 28 | G | 874 | 2 | Brahma-associated protein 60 isoform X2 | NT | NT | NT |
| 29 | P61 | 79 | 1.1 | LOC141854629 Uncharacterized protein | NT | NT | NT |
| 30 | P61 | 80 | 1 | | NT | NT | NT |
| 31 | P61 | 106 | 1.2 | Tripartite motif-containing protein brain tumor | NT | NT | NT |
| 32 | P61 | 185 | 1 | LOC141856943 Cytochrome c oxidase subunit 1-like | NT | NT | NT |
| 33 | P61 | 463 | 0.8 | | NT | NT | NT |
| 34 | P61 | 223 | 1.1 | LOC141856023 Sepiapterin reductase-like | NT | NT | NT |
| 35 | P61 | 238 | 0.9 | Uncharacterized protein LOC141857304 | NT | NT | NT |
| 36 | P61 | 320 | 0.9 | Brevipalpus obovatus Protein kinase, cAMP-dependent, catalytic subunit 1 (Pka-C1), transcript variant X1 | NT | NT | NT |
| 37 | P61 | 335 | 1.1 | LOC141853897 SREBP regulating gene protein | NT | NT | NT |
| 38 | P61 | 417 | 1.7 | LOC141849018 Uncharacterized protein | NT | NT | NT |

| **N°** | **Interactor** | **E. coli plasmid #** | **Insert size-Kb** | **BLASTn and x Result Non *Brevipalpus* sequences** | **Individual MbY2H assay** | **3-AT assay** | |
|---|---|---|---|---|---|---|---|
| | | | | | | **1** | **2** |
| 1 | G | 5 | 1.6 | *Citrus sinensis* hypothetical protein (LOC102626438) | NT | NT | NT |
| 2 | G | 111 | 0.9 | *Citrus hystrix* voucher LSY13 chloroplast | NT | NT | NT |
| 3 | G | 199 | 1.3 | *Phaseolus vulgaris* bax inhibitor 1-like | NT | NT | NT |
| 4 | G | 257 | 1 | *Vigna radiata* var. radiata uncharacterized (LOC111242361) | NT | NT | NT |
| 5 | G | 568 | 1 | *Rotaria rotatoria* NADH dehydrogenase subunit 2 | NT | NT | NT |
| 6 | G | 577 | 2 | *Gracilibacillus saliphilus* collagen-like protein | NT | NT | NT |

| | | | | | | | |
|---|---|---|---|---|---|---|---|
| 7 | G | 652 | 1 | *Penicillium brasilianum* Inorganic phosphate transport protein PHO88 | NT | NT | NT |
| 8 | G | 721 | 1 | *Philodina citrina* ATP synthase F0 subunit 6 | NT | NT | NT |
| 9 | G | 768 | 0.9 | *Neocloeon triangulifer* nuclear protein 1 | NT | NT | NT |
| 10 | G | 790 | 0.6 | *Phaseolus vulgaris* hypothetical protein PHAVU_006G185900g | NT | NT | NT |
| 11 | P61 | 182 | 1.3 | *Phaseolus vulgaris* E3 ubiquitin-protein ligase RMA1H1-like | NT | NT | NT |

| **N°** | **Interactor** | **E. coli plasmid #** | **Insert size-Kb** | **BLASTn and x Result Not hits** | **Individual MbY2H assay** | **3-AT assay** | |
|---|---|---|---|---|---|---|---|
| | | | | | | **1** | **2** |
| 1 | G | 107 | 1 | No hits | NT | NT | NT |
| 2 | G | 116 | 1.2 | No hits | NT | NT | NT |
| 3 | G | 131 | 0.8 | No hits | NT | NT | NT |
| 4 | G | 135 | 1 | No hits | NT | NT | NT |
| 5 | G | 145 | 1 | No hits | NT | NT | NT |
| 6 | G | 228 | 0.9 | No hits | NT | NT | NT |
| 7 | G | 278 | 1.4 | No hits | NT | NT | NT |
| 8 | G | 317 | 1 | No hits | NT | NT | NT |
| 9 | G | 364 | 0.7 | No hits | NT | NT | NT |
| 10 | G | 391 | 2.1 | No hits | NT | NT | NT |
| 11 | G | 418 | 1 | No hits | NT | NT | NT |
| 12 | G | 427 | 1.4 | No hits | NT | NT | NT |
| 13 | G | 558 | 1.2 | No hits | NT | NT | NT |
| 14 | G | 601 | 0.9 | No hits | NT | NT | NT |

| | | | | | | | |
|---|---|---|---|---|---|---|---|
| 15 | G | 648 | 1 | No hits | NT | NT | NT |
| 16 | G | 667 | 0.8 | No hits | NT | NT | NT |
| 17 | G | 742 | 1 | No hits | NT | NT | NT |
| 18 | G | 751 | 2 | No hits | NT | NT | NT |
| 19 | G | 895 | 0.7 | No hits | NT | NT | NT |
| 20 | G | 899 | 1 | No hits | NT | NT | NT |
| 21 | P61 | 61 | 1 | No hits | NT | NT | NT |
| 22 | P61 | 65 | 1 | No hits | NT | NT | NT |
| 23 | P61 | 284 | 0.8 | No hits | NT | NT | NT |
| 24 | P61 | 310 | 0.8-1 | No hits | NT | NT | NT |
| 25 | P61 | 357 | 0.9 | No hits | NT | NT | NT |

[1]NT: not tested; +: positive.

**Supplementary Table S3.** Summary of structural and interaction predictions for dichorhavirus and cilevirus glycoproteins and their *Brevipalpus yothersi* interactors generated using AlphaFold 3.0.

| Protein name | Organism | Size (aa)[1] | ipTM | pTM | PTM predicted | PTM type and position |
|---|---|---|---|---|---|---|
| ARF1 | *Brevipalpus yothersi* | 181 | 0.85 | 0.89 | 2 | N-Acetyl-beta-D-glucosamine (60N)<br>Phosphoserine (147S) |
| SERP2 | | 41 | 0.68 | 0.47 | 2 | N-Acetyl-beta-D-glucosamine (16N)<br>Phosphoserine (36S) |
| bryot216g00140 | | 111 | - | 0.46 | 3 | Phosphoserine (6S)<br>Phosphoserine (25S)<br>Phosphoserine (62S) |
| G | ClCSV | 491 | 0.78 | 0.6 | 12 | Phosphoserine (32S)<br>Phosphoserine (154S)<br>Phosphoserine (238S)<br>Phosphoserine (427S)<br>Phosphoserine (431S)<br>Phosphoserine (477S) |

| | | | | | | |
|---|---|---|---|---|---|---|
| | | | | | | Phosphoserine (480S)<br>Phosphoserine (483S)<br>N-Acetyl-beta-D-glucosamine (212N)<br>N-Acetyl-beta-D-glucosamine (332N)<br>N-Acetyl-beta-D-glucosamine (339N)<br>N-Acetyl-beta-D-glucosamine (368N) |
| P61 | CiLV-C | 476 | 0.51 | 0.34 | 6 | Phosphoserine (13S)<br>Phosphoserine (219S)<br>Phosphothreonine (349T)<br>Phosphoserine (365S)<br>N-Acetyl-beta-D-glucosamine (7N)<br>N-Acetyl-beta-D-glucosamine (308N) |
| **Protein-protein interaction prediction** | | | | | | |
| ARF1 x P61 | - | - | 0.16 | 0.32 | - | - |
| ARF1 x G | - | - | 0.25 | 0.34 | - | - |
| SERP2 x P61 | - | - | 0.3 | 0.38 | - | - |
| SERP2 x G | - | - | 0.45 | 0.4 | - | - |
| bryot216g00140 x P61 | - | - | 0.24 | 0.33 | - | - |
| bryot216g00140 x G | - | - | 0.37 | 0.47 | - | - |
